# Reconstituted branched actin networks sense and generate micron-scale membrane curvature

**DOI:** 10.1101/2022.08.31.505969

**Authors:** Lucia Baldauf, Felix Frey, Marcos Arribas Perez, Timon Idema, Gijsje H. Koenderink

## Abstract

The actin cortex is a complex cytoskeletal machinery which drives and responds to changes in cell shape. It must generate or adapt to plasma membrane curvature to facilitate diverse functions such as cell division, migration and phagocytosis. Due to the complex molecular makeup of the actin cortex, it remains unclear whether actin networks are inherently able to sense and generate membrane curvature, or whether they rely on their diverse binding partners to accomplish this. Here, we show that curvature sensing and generation is an inherent capability of branched actin networks nucleated by Arp2/3 and VCA. We develop a robust method to encapsulate actin inside giant unilamellar vesicles (GUVs) and assemble an actin cortex at the inner surface of the GUV membrane. We show that actin forms a uniform and thin cortical layer when present at high concentration and distinct patches that generate negative membrane curvature at low concentration. Serendipitously, we find that the GUV production method also produces dumbbell-shaped GUVs, which we explain using mathematical modelling in terms of membrane hemifusion of nested GUVs. We find that dendritic actin networks preferentially assemble at the neck of the dumbbells, which possess a micron-range convex curvature that matches the curvature generated by actin patches in spherical GUVs. Minimal dendritic actin networks can thus both generate and sense membrane curvatures, which may help mammalian cells to robustly recruit actin to curved membranes in order to facilitate diverse cellular functions such as cytokinesis and migration.

**SIGNIFICANCE:** Animal cells move, deform and divide using their actin cortex, a thin layer of filamentous proteins that supports the plasma membrane. For these actions, actin must often assemble at curved sections of the membrane, which is widely believed to require the action of dedicated actin- or membrane-bending proteins. Here, we use a bottom-up reconstitution approach to ask whether actin networks are intrinsically able to generate and sense membrane curvature. We show that membrane-nucleated actin cortices can indeed preferentially self-assemble at concave membranes generated by hemifusion of lipid vesicles. This raises intriguing questions about how such curvature recognition works, and whether cells exploit this intrinsic capability of branched actin networks to concentrate actin in specific cortical regions.

## INTRODUCTION

The actin cortex is a thin but dense meshwork of cytoskeletal actin filaments that lines the plasma membrane of animal cells. It provides mechanical stability to the cell surface, yet also actively drives cellular shape changes necessary for cell division, cell migration, and tissue morphogenesis. The main structural component of the cortex is filamentous actin. More than 100 actin-binding proteins, including nucleators, membrane-anchoring proteins, crosslinkers and motors control the organization, turnover dynamics, thickness, and mechanical properties of the cortical actin meshwork (1). Proteomic and functional studies have shown that cortical actin filaments are nucleated primarily by the formin Dia1 and the Arp2/3 complex (2, 3). Dia1 generates linear actin filaments and associates with their fast-growing (barbed) end (4), whereas the Arp2/3 complex binds to the side of a pre-existing actin filament and nucleates a new actin filament from its slow-growing (pointed) end upon activation by a nucleation promoting factor such as N-WASP (5). As a consequence of this mechanism, Arp2/3-mediated actin network growth is autocatalytic and results in dendritic arrays (6–8).

Cortical actin networks accumulate at various curved plasma membrane structures, for instance in the cytokinetic ring (9, 10), in phagocytic cups (11) and near the nucleus during cell migration in tight spaces (12). However, it remains unknown whether cortical actin networks have any intrinsic ability to sense and respond to membrane curvature. Any such abilities are difficult to observe in living cells, as cortical actin interacts with a multitude of curvature-sensitive actin-binding partners, including membrane-bending BAR-domain proteins (13–16), curvature-sensitive septins (17, 18), and IQGAP proteins (19), possibly obscuring any intrinsic curvature sensitivity of the actin network itself. Conversely, it is also unknown how cortical actin affects the generation of membrane curvature. On the one hand, polymerizing actin can generate membrane curvature by exerting pushing forces (20, 21), but on the other hand the high rigidity of the cortex may counteract membrane deformation (22, 23).

To study how simple actin networks intrinsically interact with curved membranes, we pursue a bottom-up reconstitution approach. Reconstitution of actin inside cell-sized, deformable membrane compartments has been gaining traction recently, with several studies exploring how actin networks self-organize in and shape giant unilamellar vesicles (GUVs) (20, 24–31). However, systematic *in vitro* characterization of the interplay between the membrane and a dynamic actin cortex in GUV assays has been challenging due to difficulties with reproducible actin encapsulation.

In principle, various approaches are available for GUV formation, ranging from methods based on swelling dried lipid films (32–39) through picoinjection of droplet-templated GUVs (40, 41) to emulsion-based approaches (42–51). However, most of these approaches are incompatible with actin encapsulation, as their yield drops dramatically when encapsulating proteins in physiological buffers. The most successful strategies for reconstituting actin in GUVs have been emulsion-transfer methods, where droplets of protein-containing aqueous solution are formed in an oil phase, stabilized by a self-assembled lipid monolayer, and transformed into GUVs by passing through a second lipid monolayer. One version of this type of GUV preparation is ‘continuous Droplet Interface Crossing Encapsulation’ (cDICE), a method where droplets are created by injecting a continuous stream of inner protein containing solution into the oil phase by means of a thin glass capillary. The inner aqueous phase is supplemented with a density medium, and the oil phase is atop an aqueous subphase inside a rotating chamber, so GUVs can be created on a single compact device (47). While cDICE has been used for actin encapsulation in several studies (20, 24, 25, 28, 30, 52–54), recent optimization efforts have made it significantly more robust (55), and attempts at simplifying the method have also been made (51). Nonetheless, the technique remains relatively slow and cumbersome, and a full characterization of the produced GUVs is lacking.

Here, we establish a robust and fast variant of the Cdice method which we term “emulsion Droplet Interface Crossing Encapsulation” (eDICE) to reconstitute biomimetic actin cortices inside GUVs, and characterize the resulting GUVs. We show that Arp2/3-mediated assembly generates either cortical patches or continuous cortices, depending on the concentration of actin and its nucleation factors. Interestingly, these cortical patches can generate membrane curvature. Serendipitously, we discovered that the cortical patches also recognize membrane curvature because our GUV formation protocol produces a small fraction of dumbbell-shaped GUVs that likely originate from nested GUVs. We use these dumbbell-shaped GUVs as a model system in which the membrane provides a substrate with fixed and spatially varying curvature, and find that actin networks have a strong preference for assembling at regions of concave membrane curvatures around *κ* = − 0.5 μm^− 1^.

## MATERIALS AND METHODS

### Materials

#### Chemicals

The following chemicals were purchased from Sigma-Aldrich: Tris(hydroxy-methyl)aminomethane hydrochloride (Tris-HCl), potassium chloride (KCl), calcium chloride (CaCl_2_), magnesium chloride (MgCl_2_), Optiprep (Cat. # D1556-250ML), D-(+)-glucose, DL-dithiothreitol (DTT), Adenosine 5’-triphosphate magnesium salt (MgATP), Adenosine 5’-triphosphate disodium salt (Na_2_ATP), proto-catechuic acid (PCA), proto-catechuate-3,4-dioxygenase (PCD), 3-(N-morpholino)propanesulfonic acid (MOPS), ethylene glycol-bis(β-aminoethyl ether)-tetraacetic acid (EGTA), ethylenediaminetetraacetic acid (EDTA), glycerol, β-casein, silicone oil (5 cSt), light mineral oil (BioReagent), and chloroform (Uvasol). n-decane (99% pure) was purchased from Arcos Organics. Chloroform, mineral oil and silicone oil were stored in a glove box with an ambient humidity below 1%.

#### Lipids

All lipids were purchased from Avanti Polar Lipids: 1,2-dioleoyl-sn-glycero-3-phosphocholine (DOPC), 1,2-di-stearoyl-sn-glycero-3-phosphoethanolamine-N-[methoxy (poly-ethylene glycol)-2000] (DOPE-PEG2000), 1,2-dioleoyl-sn-glycero-3-[(N-(5-amino-1-carboxypentyl) iminodiacetic acid) succinyl] (DGS-NTA(Ni)), 1,2-dioleoyl-sn-glycero-3-phospho-ethanolamine-N-(Cyanine 5) (DOPE-Cy5). DOPC was purchased in powder form and dissolved in chloroform to 25 mg/mL, while the other lipids were purchased as a solution in chloroform and used as delivered. All lipids were stored in chloroform under argon at −20° C.

#### Proteins

Rabbit skeletal muscle actin was purchased from Hypermol (Cat. # 8101-03). The lyophilized powder was dissolved according to the supplier’s instructions, resulting in a solution of monomeric (G-)actin at a concentration of 23.8 μM (= 1 mg/mL) in G-actin storage buffer (2 mM Tris-Cl, pH 8.2, 0.4 mM ATP, 0.1 mM DTT, 0.08 mM CaCl_2_ and 0.2% unspecified disaccharides). The G-actin solution was left to rest on ice for 2 hours and cleared by centrifugation at 148,000 g for 1 hour. Clearing did not result in a measurable loss of protein concentration, as verified by UV-VIS absorption measurements at 290 nm using a Nanodrop 2000c spectrophotometer using an extinction coefficient *ϵ*_290_ = 0.617^mg^/_mL_(56). Fluorescent G-actin was prepared by labelling G-actin with AlexaFluor 488 carboxylic acid succimidyl ester (Invitrogen, through Thermo Fisher Scientific, Cat. # A20000) following Ref. (57). Free dye was removed by size exclusion chromatography using a PD MiniTrap G-25 desalting column (Sigma Aldrich, Cat. # GE28-9180-07). The 10xHis-tagged VCA-domain of murine N-WASP (amino acids 400-501) was expressed in E. coli BL21 (DE3) cells and purified following Ref. (58). It was fluorescently labeled with AlexaFluor C5 maleimide (Molecular Probes) following the supplier’s protocol. Excess dye was removed from the protein by buffer exchange on a PD MiniTrap gravity column with Sephadex G-25 resin. The plasmid was a kind gift from Kristina Ganzinger (AMOLF). Human Arp2/3 isoforms ArpC1B/C5L and ArpC1A/C5 purified from SF21 insect cells was kindly provided by Michael Way (Crick Institute). Arp2/3 was stored at a stock concentration of 1 μM in Arp2/3 storage buffer (20 mM MOPS pH 7.0, 100 mM KCl, 2 mM MgCl_2_, 5 mM EGTA, 1 mM EDTA, 0.5 mM DTT, 0.2 mM ATP, 5% (v/v) glycerol). Arp2/3 protein complex from porcine brain was purchased from Hypermol EK (Cat. # 8413-01). The lyophilized protein was dissolved following the supplier’s instructions and stored at 2.23 μM in Arp2/3 storage buffer. SNAP-tagged murine capping protein (CP) was expressed in E. coli BL21 Codon Plus (DE3)-RP (Agilent Technologies) cells and purified according to Ref. (59). The plasmid was a kind gift from David Kovar (University of Chicago). After elution from Talon cobalt affinity resin, the protein was dialyzed overnight at 4° C (Cat. # D9527-100FT, 14 kDa MWCO cellulose membrane, Sigma Aldrich) into CP storage buffer (10 mM Tris-HCl pH 7.5, 40 mM KCl, 0.5 mM DTT). All proteins were snap-frozen and stored in small aliquots at −80° C. G-actin aliquots were thawed quickly and stored on ice for no more than one week. Nonlabeled and labeled G-actin monomers were mixed in a 10:1 molar ratio for fluorescence imaging. G-actin solutions were left on ice for at least 2 hours before use, to allow for depolymerization of any filamentous (F-)actin (60). Thawed aliquots of VCA, Ar2/3 and capping protein were kept on ice and used within 2 days.

### GUV preparation

#### Buffers

Buffer conditions were kept consistent between all experiments. Unless otherwise specified, the inner aqueous solution (IAS) inside the GUVs contained F-buffer (20 mM Tris-HCl pH 7.4, 50 mM KCl, 2 mM MgCl_2_, 1 mM DTT, 0.5 mM MgATP) supplemented with 0.5 μM PCD and 10 mM PCA to minimize photobleaching (61), as well as 6.5 % V/V Optiprep to increase the mass density. We reduced the Optiprep concentration as compared to the cDICE method (55) as we found that it substantially impacts actin polymerization kinetics while eDICE production of GUVs is effective over a range of Optiprep concentrations (Fig. S12). The IAS osmolarity was 168 mOsm/kg in all experiments and was unchanged by the addition of proteins, as measured using a freezing point osmometer (Osmomat 010, Gonotec, Germany).

The outer aqueous solution (OAS), in which the GUVs were first produced, contained 190 mM glucose (200 mOsm/kg). Immediately after GUV formation, we added a solution of 40 mM Tris-HCl pH 7.4 and 90 mM glucose (182 mOsm/kg), to stabilize the pH outside the GUVs and reach final buffer conditions of 10 mM Tris-HCl pH 7.4 and 170 mM glucose, with an osmolarity of 189 mOsm/kg to slightly deflate the GUVs and increase the available excess membrane area by an average of 7.6 % (62).

#### Lipid-in-oil solution

Lipid stocks were prepared in chloroform solution. 94.985 % DOPC was mixed with 0.01 % DOPE-PEG2000 to increase GUV yield following (55), 5 % DGS-NTA(Ni) to recruit VCA, and 0.005 % DOPE-Cy5 for fluorescent visualization (molar percentages). Mixtures with a total lipid content of 1.7 μmol were dried under a gentle N_2_ stream in a screw cap glass vial for each experiment and used within a day. This yields a final lipid concentration of ∼ 0.25 mmol/L in the oil phase. The vials with dried lipid films were then transferred into a glovebox under an inert environment containing <1 % environmental humidity, to promote a reproducible and high yield of GUVs (55). In the glovebox, silicone oil and mineral oil were combined in a volumetric ratio of 5.3:1.2 and thoroughly mixed by vortexing. Dried lipid films were dissolved either in 415 μL chloroform, or they were dissolved in 50 μL chloroform and subsequently diluted with 400 μL n-decane. Under continuous gentle vortexing, 6.5 mL of the oil mix was slowly added to each lipid vial. The vials were sealed tightly with teflon tape and parafilm, removed from the glove box, and sonicated on ice in a bath sonicator for 15 minutes. The lipid-in-oil solutions were stored on ice and used for GUV formation within 30 minutes.

#### GUV formation and handling

GUV preparation and imaging was performed at room temperature (24 ± 1 ° C). We used a home-built spinning chamber setup equipped with a spinning table and 3D-printed chamber described in (55) to prepare GUVs. We first set the spinning chamber to rotate at 2000 rpm (corresponding to a motor voltage of 14.5 V on our device), added 700 μL of OAS, and carefully layered on 5 mL of the oil solution. 1 mL of oil solution was set aside in a 2 mL Eppendorf tube. We then created droplets of IAS in the 1 mL oil solution by manual emulsification. 25 μL IAS were prepared on ice, adding G-actin last. The IAS was then pipetted into the 1 mL oil solution and emulsified by scratching the Eppendorf tube vigorously over an Eppendorf holder 14 times. This simple procedure allows us to form droplets within < 15 s. Note that a recent work has shown that droplets can also be prepared by vigorous pipetting (51). The emulsion was immediately pipetted into the spinning chamber and centrifuged for 3 minutes to form GUVs.

After GUV formation, the spinning chamber was stopped by slowly turning down the motor voltage, and excess oil was removed by careful pipetting. 233 μL of outer buffer were added to the remaining aqueous phase to stabilize the OAS pH, and the GUVs were concentrated by resting the spinning chamber at a 45° angle for 10 minutes. GUVs were retrieved using a cut-off 200 μL pipet tip to minimize shear forces during transfer. The samples were finally diluted threefold in imaging buffer (10 mM TrisHCl pH 7.4, 165 mM glucose).

### Microscopy

#### Image acquisition

GUVs were observed either in ibidi 18-well μ-slides (Cat. # 81817) or in chambers created by mounting silicone spacers (Sigma Aldrich, Cat # GBL665106) on # 1.5 coverslips (Superior Marienfeld, Cat. # 0107222). The wells were passivated by incubating with a 0.1 mg/mL β-casein solution in 10 mM Tris-HCl pH 7.4 for at least 15 min, rinsed with MilliQ water and dried under N_2_ flow. After GUV addition, we closed the chambers to prevent solvent evaporation and keep the osmotic deflation of the GUVs constant. Ibidi wells were closed with the appropriate lid, which effectively prevents evaporation over several days. Chambers made from silicone spacers were sealed from the top with a glass coverslip, affixed to the spacer with vacuum grease (Dow Corning high vacuum silicone grease, Sigma-Aldrich Cat. # Z273554).

Confocal images were acquired either on an inverted Olympus IX81 confocal spinning disk microscope equipped with 491 and 640 nm CW lasers, a 100x oil immersion objective (UPlanSApo, WD 0.13 mm, NA 1.4) and an EM-CCD Andor iXon X3 DU897 camera, or on an inverted Leica Stellaris 8 laser scanning confocal microscope equipped with a white light laser, a 63x glycerol immersion objective (HC PL APO, WD 0.3 mm, NA 1.2) and HyDS and HyDX detector operated in photon counting mode. Z-stacks were acquired with a 1 μm step height unless otherwise specified. Further imaging settings are listed in Supplemental Table 1.

#### Quantitative confocal microscopy

Quantitative confocal microscopy to determine the actin encapsulation efficiency in GUVs was performed on the Leica Stellaris 8 LSCM using the 63x glycerol immersion objective and the HyD S detector operated in counting mode. We performed all measurements in one day and in a single ibidi 18-well μ-slide to avoid artefacts from variations in coverslip thickness. To ensure that images were always taken at the same height above the coverslip surface, we first acquired a 256×256 pixel xz-image (Fig. S3 A). We subsequently acquired an image of 2048×2048 pixels 7 μm above the coverslip surface (Fig. S3 B). For each GUV sample, we acquired at least 10 fields of view, corresponding to > 1000 GUVs per condition. To convert fluorescence intensity to absolute actin concentrations, we obtained calibration data by imaging bulk F-actin solutions at a range of concentrations under identical imaging conditions (Fig. S3 H, I)

#### FRAP experiments

FRAP experiments to determine membrane continuity for dumbbell-shaped GUVs were performed on the Leica Stellaris 8 LSCM with a 63x glycerol immersion objective. GUV membranes were labeled with 0.05 % mol/mol DOPE-Cy5. We imaged the equatorial section of dumbbell GUVs at high zoom (typically 9x) and with a 512×512 pixel field of view. The white light laser was operated at 658 nm and at 8 % laser power with a pixel dwell time of 0.95 μs during acquisition of the pre- and post-bleach images. Bleaching was performed on a rectangular ROI, which encompassed the bright lobe of a dumbbell and was thus different in size and aspect ratio for each vesicle. The ROI was bleached using four laser lines (643, 648, 653 and 658 nm) operated at 100 % intensity each, to efficiently bleach the membrane within a single frame (= 249 ms). We acquired three images before bleaching the GUV at 4 fps. After bleaching, we acquired 50 frames at 4 fps, followed by another 50 frames at 2 fps, to capture membrane dynamics for 38 seconds.

Since dumbbell GUVs were in suspension and hence freely diffused during the course of imaging, we extracted line profiles for each frame individually, tracing the dumbbell membrane around first the bright and then the dim lobe, using the ImageJ ‘plot profile’ function (63). Time-dependent intensity ratios *I*_*bright*_ /*I*_*dim*_ were extracted from the profiles using a custom written python script.

### Image analysis

#### Quantitative analysis of actin encapsulation in GUVs

Quantitative confocal microscopy images were analyzed in an automated procedure. First, we located GUVs in the membrane images using the Template Matching module in the open source DisGUVery toolbox (64) (See Fig. S3 C). The images were preprocessed by smoothing with a filter size of 15 and edge enhancement with a filter size of 45. A representative image of a GUV membrane was used as a template, and matches with a matching index of above 0.4 were detected with 40 size steps ranging from 0.3 to 2 times the template size. Since membrane labeling was deliberately kept low to avoid any spectral crosstalk artefacts, we could only detect 30-60 % of GUVs in any given image but we observed no systematic bias in which GUVs were detected or missed. Next, we extracted the average actin fluorescence in the GUVs with a custom python script. We created a circular ROI of radius *R* = *d*_bb_/2, where *d*_bb_ is the size of the bounding box detected by the template matching GUV detection (Fig. S3 D,E). We then extracted the pixel intensities for the actin signal within the ROI and averaged these over the ROI for each GUV (Fig. S3 F). Resulting intensity distributions were converted to actin concentrations by means of a calibration curve determined for bulk actin networks (Fig. S3 H,I). For these bulk networks, we averaged the mean pixel intensity in at least 5 images of 2048×2048 px per condition. A proportional fit of measured actin signal vs nominal actin concentration in the form *I*_act_(*c*_act_) = *A* · *c*_act_ was computed using numpy least squares fitting, and the resulting proportionality constant was used to convert actin intensity in GUVs into actin concentrations.

#### Classification of actin cortex phenotype

Cortex pheno-types for each GUV were categorized manually from confocal z-stacks in one of four categories: *‘Inside patch’:* GUV has one or more actin patches in the lumen. *‘Concave patch’:* GUV has one or more actin patches which visibly protrude into the GUV lumen and/or colocalize with a section of membrane which is locally bent inwards. *‘Flat patch’:* GUV has one or more actin patches on the membrane, which do not colocalize to a concave actin patch and do not encompass the entire GUV. *‘Continuous cortex’:* GUV has an enhanced actin signal on the membrane compared to the lumen throughout the membrane. Dumbbell-shaped GUVs were excluded from the analysis of all GUVs and considered separately (below).

#### Dumbbell GUV shape analysis

The size and shape of dumbbell GUVs were extracted manually from confocal z-stacks using ImageJ (63). We fitted circular ROIs to each lobe in the equatorial slice of the dumbbell and measured the lobe diameters. We then drew a line profile across the neck of the dumbbell and extracted the distance between the two peaks in membrane signal on either side of the neck opening. Geometric features of the dumbbells (bright-to-dark size ratio, average lobe diameter, neck-to-lobe ratio) were computed from the lobe sizes and neck diameters in python.

#### Classification of actin localization on dumbbell GUVs

Actin localization in dumbbell necks was classified manually in con-focal z-stacks acquired on the Stellaris 8 LCSM. The dumbbell vesicles were first visually inspected to assess whether there was visible local actin enrichment anywhere on the membrane. It was further noted whether enrichment occured in the neck region. If so, a maximum intensity projection of the GUV was generated in ImageJ (63). On the basis of visual inspection of the projection, we classified the GUV as possessing one or more than one actin patch at the neck (*single patch* or *multi patch*), or enrichment in the entire neck region (*continuous enrichment*).

#### Quantification of membrane curvatures

Membrane curvatures in concave actin patches or around dumbbell necks were assessed using the ImageJ plugin Kappa - Curvature analysis (version 2.0.0) (65). For each patch, we drew a 5 px wide ROI around the relevant membrane section (typically 5-10 μm long), and used Kappa to extract the membrane curvature in this region, based on B-splines fit to the membrane contour. Finally, the full curves were exported to a custom python script, to compute the signed curvature along the neck.

#### Quantification of actin localization

Actin localization at membrane necks was quantified using fluorescence line profiles extracted in ImageJ. We compared intensities in two line profiles, one through the region of interest (i.e., across the dumbbell neck) and one reference region (i.e., a line along the rotational symmetry axis of the dumbbell). Line profiles of 8 px width were extracted in ImageJ and exported to a custom python script to determine the degree of enrichment as the ratio between the maximum actin intensity in the neck region, divided by the maximum actin intensity in the reference region.

## RESULTS

### A robust protocol for actin encapsulation

To study the interactions between cortical actin and lipid bilayer membranes, we reconstituted membrane-anchored actin cortices inside giant unilamellar vesicles (GUVs). We used a modified emulsion transfer method adapted from a recently optimized cDICE procedure (55) (Fig. 1 A). We used the same optimized spinning chamber setup reported in (55) and took care to maintain low humidity conditions during the procedure to enhance the GUV yield. However, rather than forming the droplets by injection of the Inner Aqueous Solution (IAS) through a capillary, we pipetted pre-formed emulsion droplets into the spinning chamber. Using pre-formed droplets has several important advantages. First, it avoids problems that can arise from clogging of the capillary needed to produce droplets with cDICE. Second, it speeds up the encapsulation step, reducing the time during which the G-actin monomers are at room temperature from several minutes to under 15 s, hence preventing premature actin polymerization (66). Third, it allows downscaling of the amount of IAS required to generate samples from > 100 μL for cDICE to < 30 μL in the case of eDICE, important when working with recombinant proteins available only in small amounts. eDICE robustly produced a high number of GUVs that are polydisperse in size and in encapsulated protein concentration. In addition to the GUVs, of which a small fraction typically had one or more small lipid pockets attached, we also usually found some large lipid aggregates (bright cyan spots in Fig. 1 B) that were not physically attached to GUVs. The average GUV size (9 *μ*m) was comparable to that obtained with optimized cDICE (12 μm (55)).

**Figure 1:**
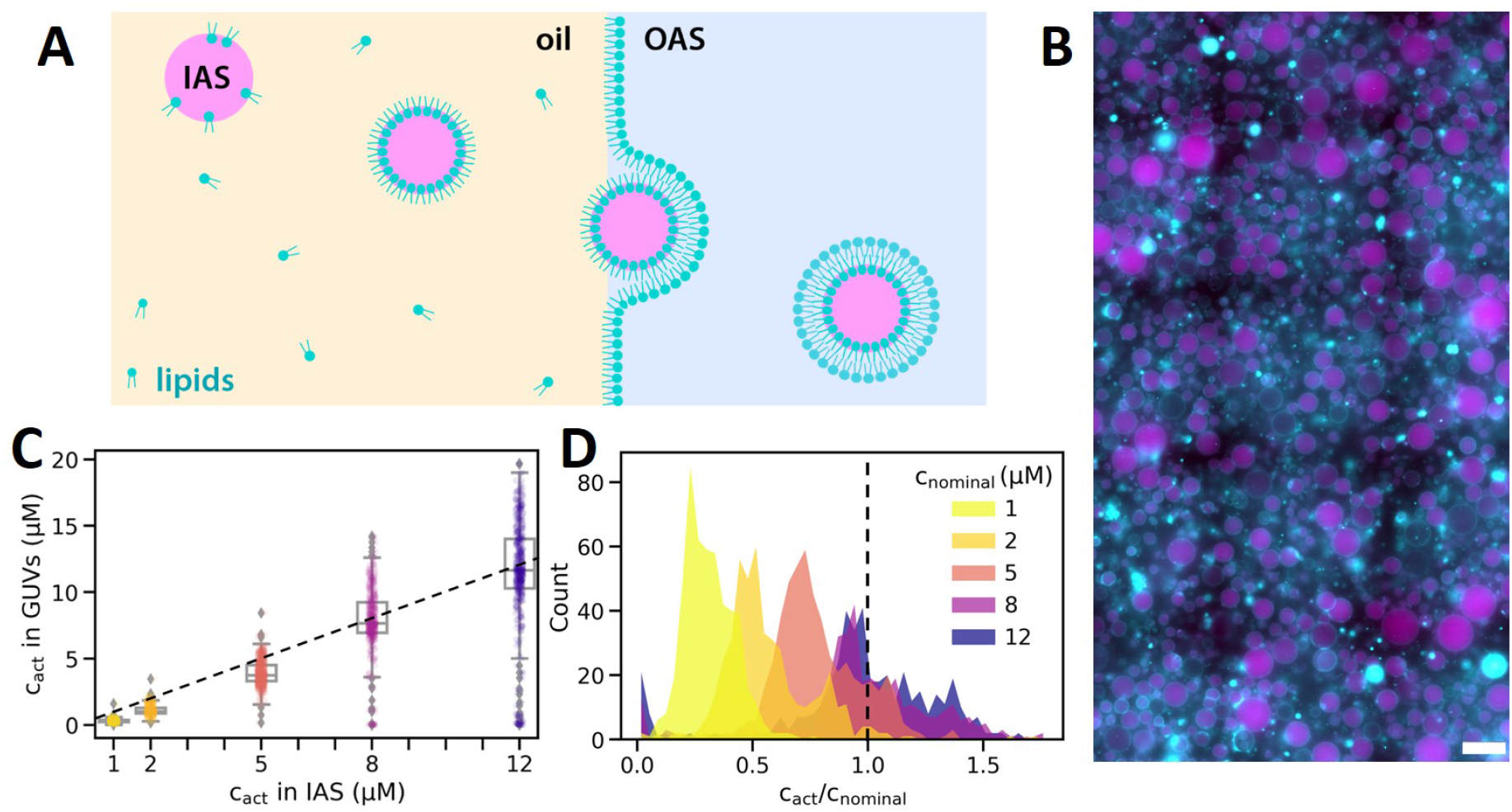
Robust actin encapsulation by eDICE. (A) Schematic of the eDICE process. Preformed lipid-stabilized emulsion droplets containing the Inner Aqueous Solution (IAS) with G-actin are injected into a spinning chamber. Centrifugal forces created by rotation of the spinning chamber establish concentric layers of the oil phase and an Outer Aqueous Solution (OAS) and push the droplets through the oil/OAS interface, where they acquire a second lipid leaflet and thus transform into GUVs. (B) Typical widefield image of GUVs formed by eDICE. F-actin is shown in magenta, lipids in cyan. Scale bar: 50 μm. (C) Actin concentrations in eDICE GUVs measured by quantitative fluorescence microscopy, as a function of the nominal actin concentration in the IAS. Individual data points indicate single GUVs, boxplots indicate medians and quartiles. (D) Histogram of encapsulation efficiency, *c*_rel_ = *c*_act_/*c*_nominal_, for different nominal actin concentrations. The dashed line represents *c*_rel_ = 1. *N*_*c*_ = 515, 468, 536, 368 and 447 GUVs for *c* = 1, 2, 5, 8 and 12 μM actin, respectively.

To quantify the encapsulation efficiency, we analyzed confocal fluorescence images of actin inside the GUVs and converted intensities to concentrations using calibration measurements on bulk actin solutions. We found that the mean actin concentration inside the GUVs increased linearly with the concentration of actin in the IAS, and that the mean value was close to the nominal (input) actin concentration (Fig. 1 C). However, the distribution of actin concentrations broadened as the nominal actin concentration increased. Interestingly, at high actin concentrations (*c*_nominal_ ≥ 8 μM), a substantial fraction of the GUVs was supersaturated with actin, containing up to 1.7× the nominal actin concentration, while a small fraction of the GUVs showed no measurable actin content at all (Fig. 1 D). Extended quantification of these results is shown in Fig. S4. At low concentrations of actin in the IAS (*c*_nominal_ ≤ 2 μM)), a large fraction of GUVs contained substantially less actin than expected (Fig. 1 D). While it is widely known that GUV formation yields a polydispersity in solute concentrations, our results highlight that this polydis-persity is in fact nontrivially dependent on the initial solute concentration.

### Actin cortex formation proceeds by nucleation and fusion of patches

In order to form a membrane-nucleated actin cortex, we encapsulated actin along with the nucleating protein Arp2/3 and the Arp2/3-activating protein VCA (Fig. 2 A). Nucleation was constrained to the membrane by using a 10×-His tag on the VCA protein in order to recruit it to the GUV membrane via Ni-NTA lipids present at a small density within the membrane.

**Figure 2:**
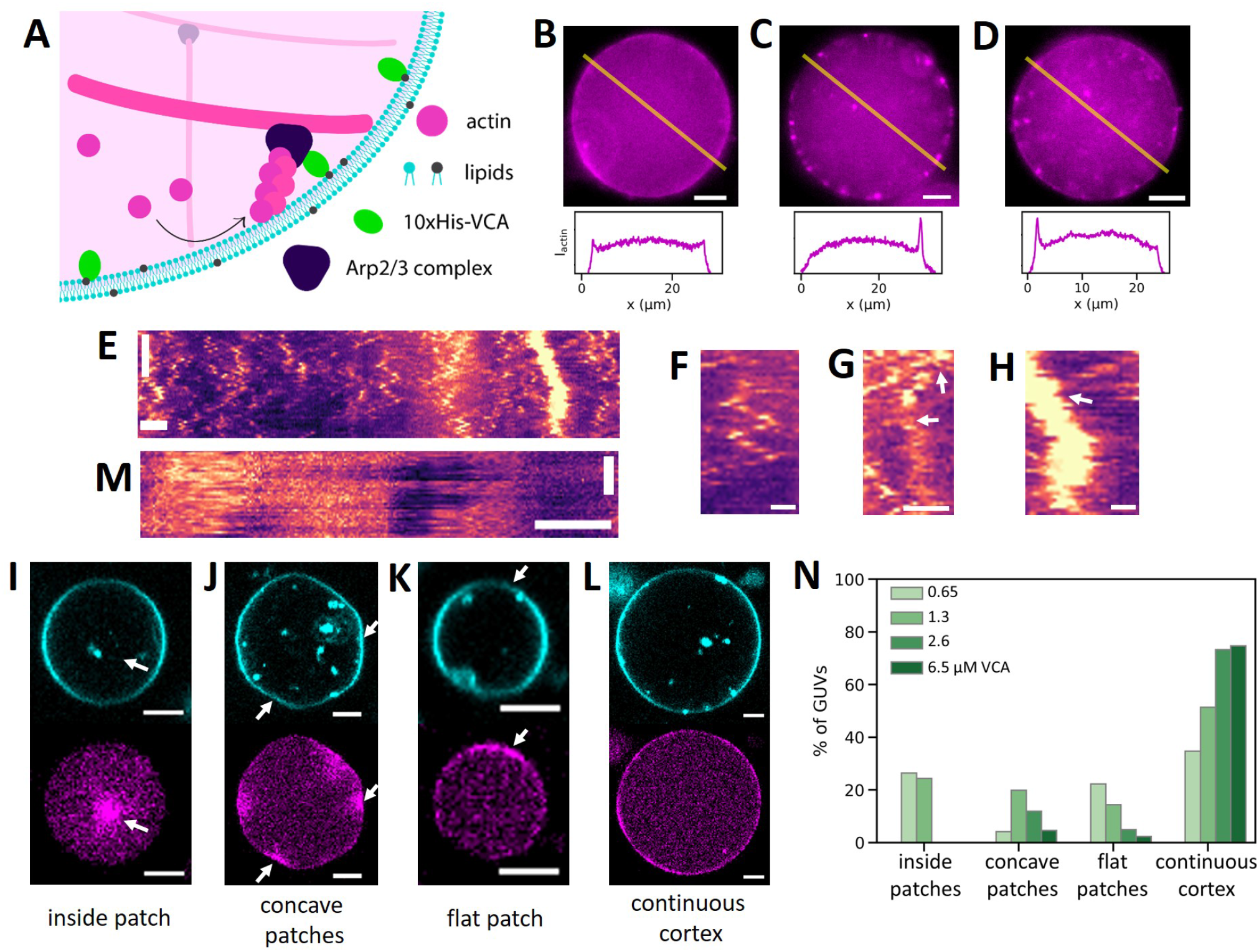
Formation of membrane-nucleated actin cortices inside GUVs. (A) Actin filaments (magenta) were nucleated using the Arp2/3 complex, which was activated near the inner leaflet of the GUV membrane (cyan) by membrane-bound VCA (green). (B - D) Top row: At low actin concentration (4.4 μM), actin formed either a continuous cortex (B), or small bright patches (C), or a combination of both (D). Bottom row: Line profiles along the yellow lines in the epifluorescence images. (E) Kymograph of a patchy actin cortex from a line drawn along the circumference of a GUV in a time lapse of 50 frames recorded at a frame rate of 1 fps. Scale bars: 5 μm (horizontal), 20 s (vertical, arrow of time points downwards). (F - H) Zoomed-in sections of the kymograph shown in (E). Actin patches could diffuse along the cortex (F), split (G, white arrows) or merge (H, white arrow). Scale bars: 2 μm. (I - L) At high actin concentration (8 μM), we found bright actin foci inside the lumen (I, white arrow), bright concave actin patches (J, white arrows), extended flat membrane patches (K, white arrow), or a continuous flat cortex (L). All examples show GUVs with 2 μM VCA and 50 nM Arp2/3. (M) Kymograph of an actin cortex (8 μM actin) with large flat patches that were immobile. Scale bars: 5 μm (horizontal), 20 s (vertical). (N) Quantification of cortical phenotypes for 8 μM actin and 50 nM Arp2/3 as a function of the density of VCA activator. Actin patches in the GUV lumen occurred only at low VCA concentrations, concave patches were most prevalent at intermediate VCA concentrations, continuous cortices were most prevalent at high VCA concentrations. GUVs were all produced on the same day and statistics are based on *N* = 72, 111, 101 and 87 GUVs for VCA concentrations of 0.65, 1.3, 2.6 and 6.5 μM, respectively.

At low actin concentrations (4.4 μM), actin cortices formed with 50 nM Arp2/3 and 0.65 μM VCA exhibited three distinct types of structures. Cortices were either continuous, being enriched all around the GUV (Fig. 2 B), or they were made up of distinct small (1-2 μm-sized) cortical actin patches (Fig. 2 C), or they showed a weakly enriched signal throughout the cortex as well as enrichment in separate patches (Fig. 2 D). Kymographs of the actin signal along the membrane (Fig. 2 E) revealed that the actin patches diffuse over the surface on a timescale of seconds (Fig. 2 F) and can also split (Fig. 2 G) or merge (Fig. 2 H). We note that the prevalence of the three cortex phenotypes varied widely from day to day, likely due to variations in protein encapsulation efficiency.

When we increased the nominal actin concentration to 8 μM, observations varied much less from day to day. Furthermore, at 8 μM, we no longer found any GUVs with small cortical actin patches. Instead, we observed four different cortical phenotypes. We found GUVs with bright patches of actin in the lumen (Fig. 2 I), patches of actin that were localized to concave sections of the membrane (Fig. 2 J), large flat patches of cortical actin (Fig. 2 K), and GUVs with a continuous actin cortex (Fig. 2 L). To test the role of the nucleator density, we increased the VCA concentration, which determines both the rate of actin nucleation and the interaction strength of actin with the membrane, while maintaining a constant actin concentration of 8 μM. The phenotypes we observed remained the same across VCA concentrations ranging from 0.65 to 6.5 μM, but their prevalence changed with increasing VCA concentration. For low VCA concentrations (≲ 2 μM), actin formed bright foci in the lumen of GUVs (Fig. 2 I). At VCA concentrations above 2 μM, we no longer found any actin patches in the lumen, but only at the cortex. Across the GUV populations, we found patches inside the GUV lumen in ∼ 23 % of GUVs with 0.65 or 1.3 μM VCA and in a few cases at 2 μM (Fig. 2 I), but never at higher VCA concentrations (Fig. 2 N). The cortical patches observed at VCA concentrations above 2 μM took on three configurations: extended patches that are either concave (Fig. 2 J) or flat (Fig. 2 K), and continuous cortices (Fig. 2 L). The extended patches were immobile over a 50 s timespan (kymograph in Fig. 2 M), in contrast to the small mobile patches formed at 4.4 μM actin. Flat patches strongly declined in frequency from 22 to 2 % of GUVs when the VCA concentration was raised from 0.65 to 6.5 μM, while the fraction of GUVs with a continuous cortex rose from 35 % to 75 % over this VCA concentration range (Fig. 2 N). Since patches tend to become larger compared to the GUV perimeter as VCA concentration increases, we eventually classify them as a continuous cortex. Concave patches were observed both in the presence and absence of a continuous cortex. Their prevalence peaked at a 1.3 μM VCA concentration and dropped to around 4 % of GUVs at both ends of the studied VCA concentration range.

### eDICE produces some dumbbell-shaped GUVs due to membrane hemifusion

In all GUV samples produced by eDICE, we found some GUVs with a striking dumbbell shape (Fig. 3). The prevalence of such dumbbell-shaped GUVs varied from experiment to experiment, but they usually made up a few percent of all observed GUVs. These GUVs consisted of two spherical caps connected by an open neck (Fig. 3 A). They can be distinguished from GUV doublets, created when two vesicles stick to one another (Fig. 3 B), by the fact that they lack a membrane septum. Strikingly, one lobe of the dumbbell appeared as a normal unilamellar vesicle membrane (right lobe in Fig. 3 A), whereas the other appeared either as one bright membrane (left lobe in Fig. 3 A) or as two separate membranes (Fig. 3 C), which could be be close together (left panel) or far apart (right panel). Line intensity profiles along the membrane contours revealed that the brighter lobe showed on average twice the membrane intensity of the dimmer lobe (Fig. 3 D, E). Thus, dumbbell-shaped GUVs appeared to be made up of one lipid bilayer on the dim side and two on the bright side, consistent with the observation of dumbbells with two well-separated membranes in the bright lobe (Fig. 3 C). The intensity ratio of ∼ 2 also confirmed that the dumbbell structures could not be two vesicles stuck to one another, which would have three lipid bilayers on the bright side.

**Figure 3:**
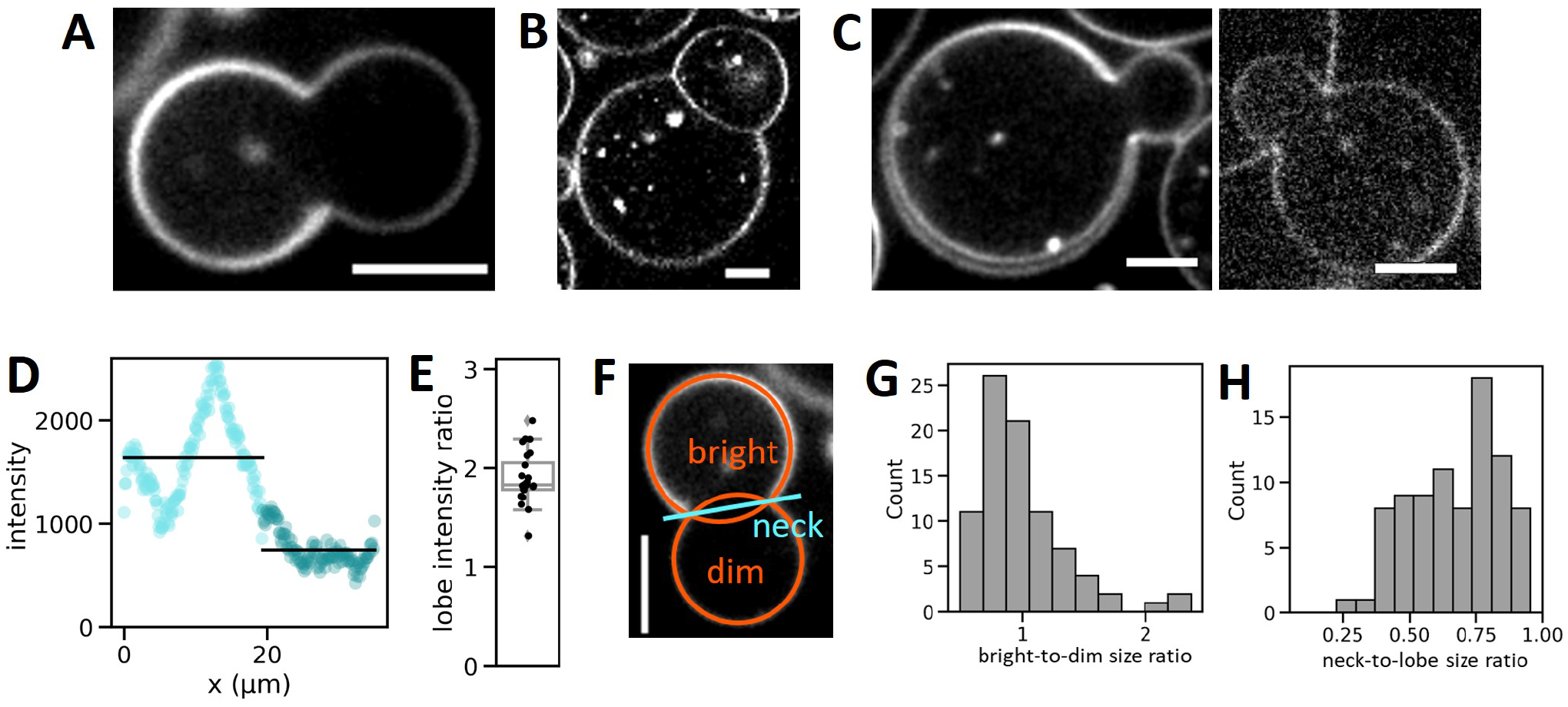
Dumbbell-shaped GUVs formed by eDICE. (A) Dumbbell-shaped GUV with a bright and a dim spherical cap connected by an open membrane neck. (B) GUV doublets clearly look different, exhibiting a membrane septum between the two lobes that each have equal membrane intensity. (C) Some dumbbells showed two clearly separate membranes in (parts of) the bright half of the dumbbell with a narrow (left) or wide (right) gap. (D) Membrane signal in a 5 pixel-wide line, starting at the neck and following clockwise along the GUV membrane shown in (A). Light cyan data points denote the bright half of the GUV, dark cyan data points show the dimmer half. The average signal is 1.8-fold higher (2.1-fold after background subtraction) in the bright than in the dim lobe (black lines). (E) Boxplot of membrane intensity ratios after background subtraction. The average ratio is 1.9 (*N* = 25 dumbbells from 5 independent experiments). (F) We measured morphological features of GUV dumbbells by fitting circles to their bright and dim lobes (orange circles) and measuring the neck diameter as the distance between the maxima in membrane fluorescence in a line profile drawn through the neck. (G) Histogram of the size ratio between bright and dim dumbbell lobes (*N* = 85). (H) Corresponding histogram of neck-to-lobe size ratios (*N* = 85). Scale bars: 5 μm.

We characterized the shapes of dumbbell vesicles by fitting circles to both lobes in the equatorial plane and measuring the distance between the two membranes at the neck (Fig. 3 F). This analysis revealed that dumbbell shapes were diverse but followed general overall trends. First, the two lobes were usually similar in size, with the dim lobe being slightly larger than the bright lobe in most GUVs (64 % of *N* = 85 GUVs), although there were occasional outliers where the bright lobe was more than twice the size of the dim lobe (Fig. 3 G, Fig. S5). The neck diameter was usually quite wide, ranging from 0.4 to 0.9 times the average lobe diameter (Fig. 3 H). Only ∼ 5 % of dumbbells had a neck narrower than 0.4 times the average lobe size.

How could these dumbbell-shaped GUVs arise? Motivated by the observation that a small fraction of GUVs contain another GUV within their lumen (Fig. 4 A), we hypothesized that dumbbells might originate from bursting of such enclosed GUVs and hemifusion with the lipid bilayer of the outside GUV (Fig. 4 B). How exactly this happens remains unclear, but we suspect that residual oil in the membrane may promote the formation of membrane defects and facilitate membrane fusion. This process should create an interface between the single and double membrane that is associated with a line tension, which is expected to drive the formation of a dumbbell shape (Fig. 4 B). Since one lobe of this dumbbell (the bright lobe) should contain one lipid double layer that is disconnected from the rest of the GUV (Fig. 4 C), we tested this interpretation by fluorescence recovery after photobleaching (FRAP). Note that this double layer really comprises only one continuous leaflet, but is bent in on itself such that it appears as two layers (illustrated in Fig. S6 C). The ratio between the average membrane intensities in the bright and the dim lobe should be 2 before bleaching, drop to 0 when we completely photobleach the bright lobe, and return to 1 over time, as the outer leaflets of the bright lobe are replenished by fluorescent lipids from the dim lobe, while the disconnected leaflets in the bright lobe remain bleached (Fig. S6). By contrast, if the leaflets in the two lobes were either fully connected or fully disconnected from one another, the final intensity ratio would be 2 or 0, respectively. Since lipids diffused rapidly in the membrane and recovered after photobleaching within less than 3 s (Fig. S7 A, B), we could not completely bleach the bright lobe to zero while retaining enough membrane intensity in the dim lobe for data analysis. We could, however, reduce the bright lobe fluorescence to around 30 % of its initial value. We found that the lobe intensity ratio recovered within a few seconds and indeed reached a final value that was just over half of its initial value (Fig. 4 D and additional data in Fig. S7 A), confirming that the lobes were hemifused.

**Figure 4:**
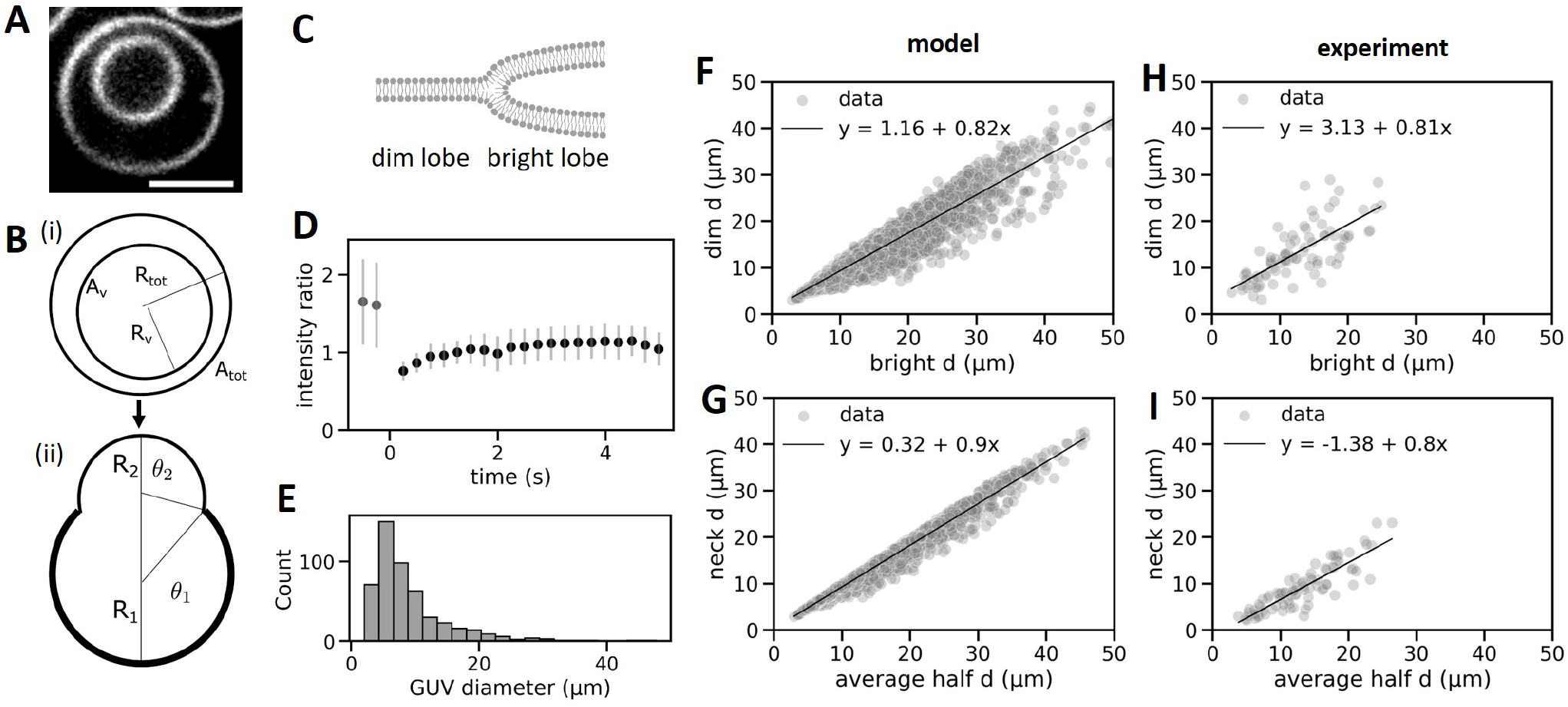
Dumbbell shapes result from hemifusion and are determined by osmotic pressure and line tension. (A) Confocal image of an eDICE GUV encapsulating another GUV (scale bar: 5 μm). Such nested GUVs were rare (∼ 5-10 % of GUVs) but present at frequencies comparable with dumbbells. (B) Proposed mechanism for dumbbell formation. (i) One vesicle with radius *R*_tot_ encapsulates another vesicle with radius *R*_v_. (ii) The inner GUV bursts and hemifuses with the membrane of the outer GUV. The shape of the resulting dumbbell is determined by membrane tension and line tension along the hemifusion line. (C) Proposed configuration of the dumbbell neck, where one inner and one outer leaflet (dim lobe, left) join four concentric leaflets in the bright lobe (right). (D) FRAP measurement of the bright lobe of a dumbbell reveals a recovery of the normalized fluorescence intensity from ∼ 2 (pre-bleach, *t* < 0) to ∼ 1 (post-bleach, *t* > 0) within seconds. (E) Experimentally measured GUV size distribution, which served as input for the statistical model. (F, G) The model predicted dumbbell shapes (relative lobe sizes, (F), and neck diameters relative to the dim lobe size, (G)) that quantitatively match the experimental data (H, I). Experimental and simulated data represent *N* = 85 and *N* = 10^4^ dumbbell GUVs, respectively. Lines and legends display linear fits.

To further test whether a model based on hemifusion of nested GUVs explains the formation of dumbbells, we asked whether the dumbbell geometries were also quantitatively consistent with the geometries expected based on a simple membrane model. We assumed that a GUV of radius *R*_tot_ encapsulates another vesicle of radius *R*_v_ (Fig. 4 B). After the encapsulated vesicle bursts, we expect its membrane to stick to that of the outer vesicle. As the membrane area of the inner vesicle is smaller than that of the outer one, this process will inevitably create two regions, one with a single and another with a double membrane. The regions will therefore have different surface tensions, and experience a line tension at their boundary. We can account for these differences in a simple energy model (see Section I of the Supplemental Material), from which we find that dumbbell shapes are indeed equilibrium solutions. Unsurprisingly, we found that the surface tensions are related to the osmotic pressure difference across the membrane according to Laplace’s law (Eqn. 5 in Supplermental Material). Moreover, we found that the overall shape of the dumbbell GUVs is the result of the interplay between this osmotic pressure, which promotes spherical shapes, and the line tension, which tends to constrict the boundary between the two lobes (Eqn. 7 in Supplermental Material). Indeed, similar dumbbell shapes have been observed and theoretically predicted in case of GUVs made of phase-separating lipid mixtures, where line tension along the domain boundary also produces spherical cap shapes (67–70).

To verify the model, we used it to predict the distribution of dumbbell shapes, based on the observed distribution of (spherical) GUV sizes. From the model, we can then calculate the ratio of the line tension to the osmotic pressure, as the physical parameter that sets the dumbbell shape. This value, which is difficult to measure directly, can thus be inferred from easily observable quantities like the dumbbell radii. We could also compare it to values for multi-component membranes, which, due to phase separation, also develop multiple domains with line tensions at their boundaries.

In the model, the total membrane area (the surface area plus excess membrane area) of the bursting vesicle *A*_v_ equals the surface area of the double bilayer membrane lobe *A*_1_, i.e. *A*_v_ = *A*_1_. Moreover, the GUV’s total membrane area *A*_tot_ minus the surface area of the bursting lobe *A*_1_ equals the surface area of the single bilayer membrane lobe *A*_2_, i.e., *A*_tot_ − *A*_1_ = *A*_2_ (Fig. 4 B (ii)). We calculated the radius of the bright lobe, *R*_1_, from the total membrane area of the bursting vesicle, *A*_v_,

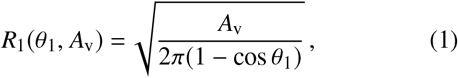

where *θ*_1_ is the opening angle of the spherical cap. Similarly, we calculated the radius of the dim lobe, *R*_2_,

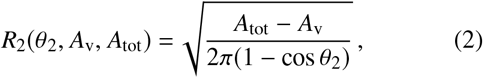

where *θ*_2_ is the opening angle of the spherical cap. The two caps are connected at a circular interface with neck radius

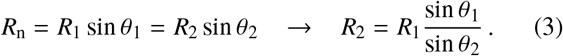

By equating Eq. (2) with Eq. (3) we obtain

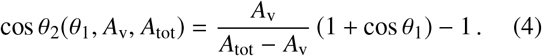

The total volume of the GUV is given by adding up the volumes of the spherical caps of the bright and dark lobe

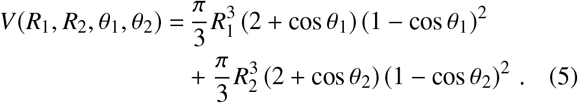

By substituting Eqs. (1), (2) and (4) in Eq. (5), we got an expression for the volume *V* = *V* (*θ*_1_, *A*_tot_, *A*_v_) of the GUV that only depends on *θ*_1_, *A*_tot_ and *A*_v_.

To connect the shape description derived above to the experiments, we calculated the geometrical correlations between the diameter of the bright and dim lobes from experimental data. First, we assumed that in the model, similar to the experiments, GUVs are created with excess membrane area that allows the GUV to change shape at constant volume. Therefore, the total membrane area *A* is larger than the surface area *A*_s_ = 4*πR*^2^, where *R* is the radius of the spherical GUV. In this case, the reduced area is given by *ν* = *A*/*A*_s_, which varies across GUVs. Next, we sampled two radii *R*_tot_, *R*_v_ from the measured diameter distribution of GUVs (Fig. 4 E) that represent the sizes of the outer vesicle and the enclosed vesicle, respectively. From the radii *R*_tot_, *R*_v_ and the reduced areas *ν*_tot_, *ν*_v_ we calculated the GUV total membrane area,

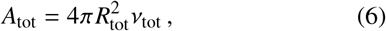

and the total membrane area of the enclosed vesicle,

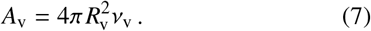

We calculated the initial (spherical) GUV volume 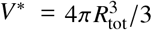 and set *V* ^∗^ equal to the dumbbell volume obtained from Eq. (5),

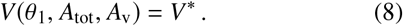

We solved Eq. (8) for *θ*_1_ by using the values of *A*_tot_ and *A*_v_ from Eqs. (6) and (7). With the value of *θ*_1_ and the values of *A*_tot_ and *A*_v_ we could calculate the remaining quantities, where we determine *θ*_2_, *R*_1_, *R*_2_ and *R*_n_ from Eqs. (4), (1), (2) and (3), respectively. Together, *θ*_1_, *θ*_2_, *R*_1_, and *R*_2_ define the GUV dumbbell shape. To determine the shapes of dumbbell GUVs, we sampled 10 000 radii *R*_tot_ and *R*_v_ and only recorded values of *R*_1_ and *R*_2_ that were in the experimental range (< 25 μm) and corresponded to a dumbbell shape (0 < *θ*_1_, *θ*_2_ < *π*). To account for statistical variability in the total membrane area of the GUV and the encapsulated vesicle, we drew the reduced areas *ν* from a truncated normal distribution with mean 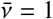 and standard deviation *σ*, where we exclude nonphysical values (*ν* < 1). While we took 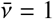 to be fixed by the initial spherical GUV shape, we treated *σ* as a free parameter. We started with *σ* = 0.01 and increased *σ* in steps of 0.01 until the slope of the linear fit of the simulated data (Fig. 4 F) matched the slope of the linear fit of the experimental data (Fig. 4 H). We found the best agreement for a value of *σ* = 0.03. For this value also the neck diameter shows a linear dependence on the average lobe diameter (Fig. 4 G).

We found remarkable agreement with linear relationships between bright and dim lobe diameters for the model and the experimental data (Figs. 4 F, H). The same holds for the relation between the neck diameters and average lobe diameters (Figs. 4 G, I). The slopes of the linear fits to the experimental and simulation data even match quantitatively, with slopes of 0.81 and 0.82 relating the dim and bright lobe diameters, and 0.8 and 0.9 relating the neck diameter and average lobe size. The abscissa of the linear fits differed by around 2.0 μm between simulated and experimentally measured relative lobe sizes (Figs. 4 G, I). This difference is likely an artefact of the way in which we select the size of the inner vesicle in the model: we choose a size from the same distribution as the outer GUV size. However, it is currently unclear how the inner vesicles actually form. It is therefore possible that the typical difference in total membrane area between inner and outer GUV is greater in the experiments than in our simulations, thus resulting in an offset between the experimental and simulated data.

Based on the good agreement of the experimentally observed shapes with a model where line tension drives dumbbell formation, we concluded that membrane hemifusion is indeed sufficient to drive the formation of dumbbell GUVs. This conclusion was supported by the observation that the same geometrical correlations arose whether or not any actin was encapsulated in the GUVs (Fig. S8 A, B). The conclusion that membrane properties alone are sufficient to form dumbbell GUVs was further confirmed by the fact that they occur in eDICE GUVs irrespective of the composition of the inner aqueous solution, as well as by the fact that we occasionally found the similar dumbbells in GUV samples produced by cDICE or gel-assisted swelling (Fig. S8 C, D).

Based on the strong agreement between the model predictions and experimental data, we now inverted the workflow and used the model to calculate the ratio between the line tension *σ* and osmotic pressure difference *P* between the inside and the outside of the vesicle, for the measured GUV dumbbell shapes. This ratio can be calculated directly from the neck radius *R*_n_ and opening angles *θ*_1_, *θ*_2_ of a dumbbell vesicle (Eq. (S7) in the Supplemental Material):

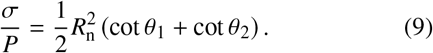

We calculated this ratio for *N* = 80 dumbbell GUVs (Fig. S2), and found a mean value of − 21 *μ*m^2^, which is around half of the measured value for a vesicle that undergoes budding due to phase separation of the membrane into different domains (69).

### Arp2/3-nucleated actin networks recognize membrane curvature

Having established that dumbbell GUVs form by a mechanism that is independent of the presence of an actin cortex, we considered them a convenient template on which to study how membrane-nucleated actin cortices react to externally imposed variations in curvature. Due to the low number of dumbbell GUVs produced in one eDICE experiment, we could not directly compare dumbbells from a single experiment with good statistics. The following discussion will thus encompass GUVs from 7 different experiments and with several slightly different cortex compositions. We mainly used the human Arp2/3 isoform C1BC5L, but we obtain similar data for human Arp2/3C1AC5 and for bovine brain Arp2/3 (Fig. S9 A).

We found four different cortical actin distribution phenotypes in dumbbell shaped GUVs. First, actin could form a continuous and uniform cortex (Fig. 5 A). Second, we found GUVs where actin was enriched in a single patch at the GUV neck (Fig. 5 B). Third, actin formed a cortex throughout but was enriched in several distinct patches along the neck (Fig. 5 C). Fourth, the entire neck could be enriched in actin compared to the rest of the cortex (Fig. 5 D). These four phenotypes occurred at roughly the same frequency (around 25 % of N=27 dumbbells, Fig. 5 E). Importantly, we found that in cases where actin formed patches, there were always patches present at the neck. Rarely, we found a GUV which had bright actin patches both at the neck and elsewhere (Fig. 5 C). Preferential actin localization to the neck did not appear to be dependent on the presence of a continuous cortex: we found a few dumbbell GUVs where the cortical region was not significantly enriched in actin compared to the GUV lumen, but also for these, around 75 % exhibited actin enrichment at the neck (Fig. 5 F, N=27 in total). The likelihood that actin was enriched at a dumbbell neck depended on VCA concentration, with 88 % of dumbbells showing neck enrichment at 2 μM VCA, compared to 68 % in dumbbells at 6.5 μM VCA (Fig. 5 G, N=8 and 19 GUVs, respectively).

**Figure 5:**
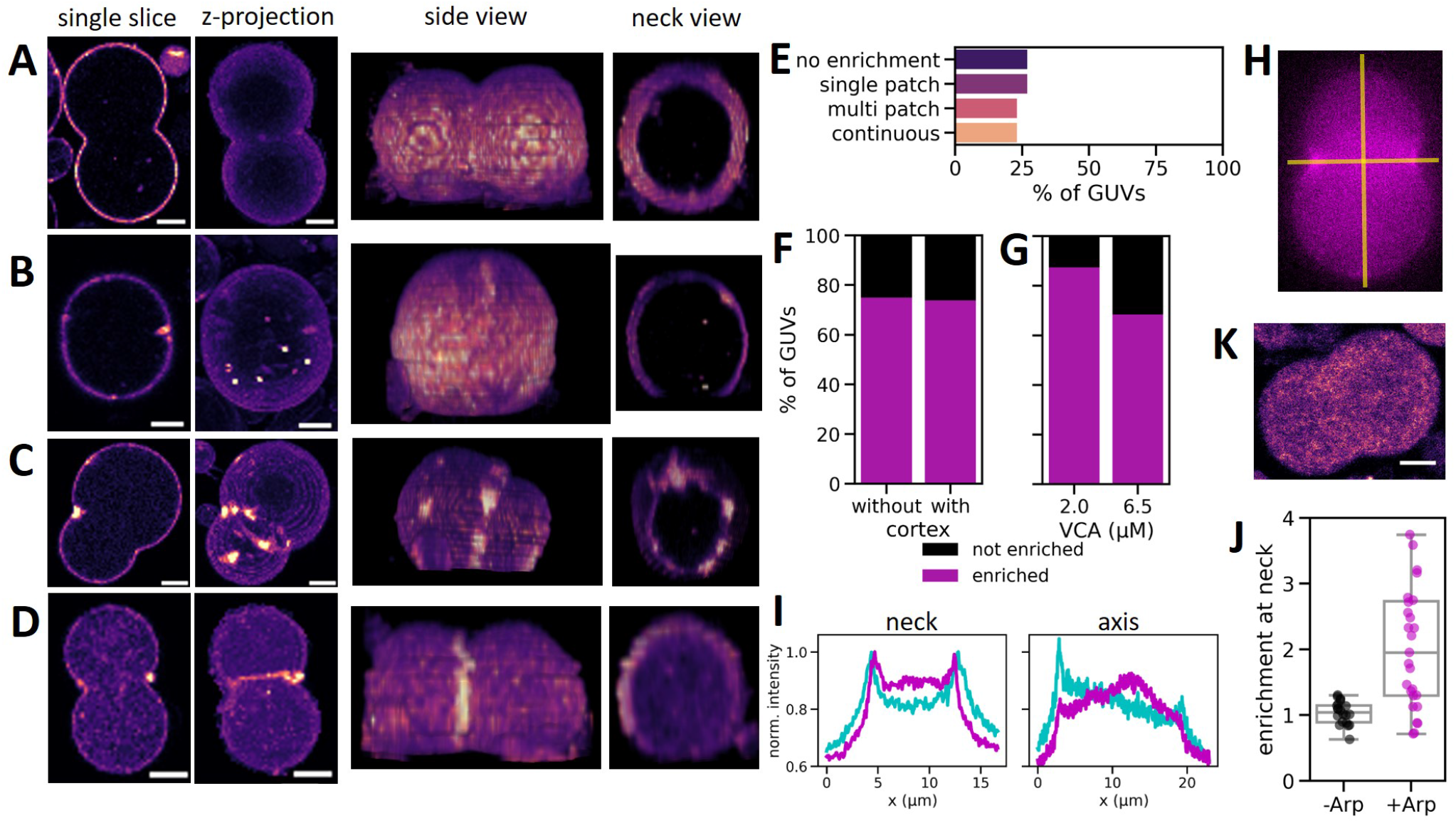
Cortical actin networks preferentially localize to dumbbell necks, indicating curvature recognition. (A-D) Dumbbell GUVs containing cortical actin showed four phenotypes: actin was evenly distributed (A), enriched in a single patch at the neck (B), enriched in several distinct patches around the neck (C) or enriched in the entire neck region (D). Columns show a single confocal slice, a maximum intensity projection, a sideview of the 3D-reconstructed z-stack, and a view through the neck section in the z-stack. Actin intensity is shown in false color (magma) for clarity. (E) Bar plot of actin enrichment patterns in dumbbell shaped GUVs, showing an even distribution over the four groups of (A-D) (N=27 dumbbell GUVs). (F) Bar plot shows that 75% of dumbbell shaped GUVs shows actin enrichment at the neck, both for GUVs with and without a continuous actin cortex (N = 23 and 4, respectively). (G) Bar plot shows that actin enrichment at the neck was slightly more common in GUVs with 2 μM VCA compared to 6.5 μM VCA (N = 8 and 19, respectively). (H) We quantified the amount of actin enrichment at the neck by comparing line intensity profiles across the neck versus the dumbbell’s symmetry axis (yellow lines). (I) Line profiles of membrane (cyan) and actin (magenta) intensity along the neck and symmetry axis of the dumbbell GUV shown in (H), normalized to the maximum pixel value in the neck profiles. Peaks in actin and membrane intensity coincide at the neck, but not at the poles. (J) Box plot of the degree of actin enrichment at dumbbell necks in GUVs where actin nucleated spontaneously in the GUV lumen (‘-Arp’, black dots, N=21) or at the membrane with the help of Arp2/3 (‘+Arp’, magenta dots, N=27). (K) Confocal image of a dumbbell where actin polymerized spontaneously in the GUV lumen and does not localize to the neck region. Scale bars: 5 μm.

We quantified the degree of actin enrichment at the neck by comparing the maximum actin intensity in a line profile through the neck with that measured in a line profile through the dumbbell’s axis of rotational symmetry (i.e., along the yellow lines in Fig. 5 H, resulting line profiles in Fig. 5 I). The degree of actin enrichment in the neck was quite widely spread, ranging from no enrichment (*I*_max, neck_/*I*_max, axis_ ≈ 1) to an almost four-fold enrichment at the neck compared to the poles (*I*_max, neck_/*I*_max, axis_ ≈ 4), with a mean enrichment of *I*_max, neck_/*I*_max, axis_ = 2.2 (Fig. 5 J, magenta points, N=27).

Importantly, in absence of actin, VCA itself did not preferentially accumulate at dumbbell necks (Fig. S9 B). Also, neck enrichment was only present in GUVs where actin was nucleated on the membrane with the help of Arp2/3 and VCA. In GUVs where actin spontaneously nucleated in the GUV lumen without these nucleating proteins, we observed no enrichment at the neck (Fig. 5 K and black points (N=21) in Fig. 5 J).

### Arp2/3-nucleated actin networks can generate membrane curvature

Our data clearly demonstrates that Arp2/3-nucleated actin networks preferentially assemble at regions of negative membrane curvature, with the dumbbell-shaped GUVs providing a template for such curvature-sensitive actin enrichment. Conversely, it has previously been reported by Dürre et al. (20) that dendritic actin networks can generate concave actin-rich domains in GUVs, which in their case required the presence of capping protein. Interestingly, we also observed concave actin-rich domains in GUVs containing 8 μM actin, but no capping protein (see example in Fig. 2 J). The concave actin patches (Fig. 6 A, see white arrowheads) co-localized with regions of negative membrane curvature (Fig. 6 B, grey shaded regions). When we quantified the prevalence of such concave patches, we found at least one concave actin patch in 37 % of GUVs, although the GUVs were more likely to have a continuous actin cortex (72 % of GUVs, see Fig. 6 C, N = 60 GUVs). When we added 185 nM capping protein, we observed a strong increase in the prevalence of concave patches, which is consistent with the study of Dürre et al. (20): we now found concave patches in 69 % of N = 36 GUVs (Fig. 6 A and C) and continuous cortices or flat patches in only 3 or 5 % of GUVs, respectively (Fig. 6 C).

**Figure 6:**
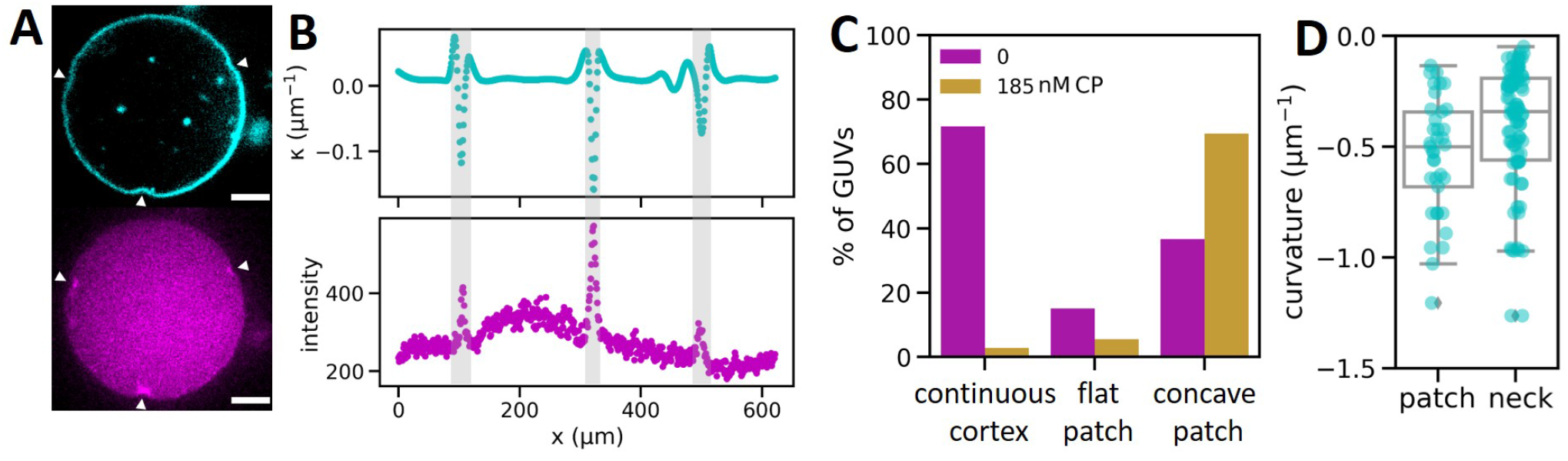
Branched actin networks both generate and sense membrane curvature. (A) Confocal slice of a GUV co-encapsulating 8 μM actin, 50 nM Arp2/3, 2.6 μM VCA and 185 nM capping protein. Actin (magenta) is enriched in small, bright, concave patches (white arrowheads) colocalized with bent sections of the GUV membrane (cyan). (B) Line profiles of the local membrane curvature (top) and actin signal intensity (bottom) along the GUV membrane in (A) confirm that spikes in actin signal occur at locations of negative membrane curvature (grey shaded regions). (C) Bar plot of the fraction of non-dumbbell GUVs with a continuous cortex, flat cortical patches, or concave cortical patches, for cortices without (magenta, N = 60 GUVs) or with (yellow, N = 36 GUVs) 185 nM capping protein. (D) Box plot of maximum negative membrane curvatures in concave actin patches in spherical GUVs (N = 37 patches in 20 separate GUVs) or at necks of dumbbell GUVs (N = 94 neck profiles from 47 dumbbells). Scale bars: 5 μm.

To assess whether the geometry of the concave actin patches formed in spherical GUVs resembles that which the cortical actin encounters at the neck of GUV dumbbells, we quantified the curvature of both (Fig. 6 C). We found that the typical curvatures of concave membrane patches (mean *κ* = 0.54 μm^− 1^, N = 37) and dumbbell necks (mean *κ* = 0.43 μm^− 1^, N=94) are indeed not statistically different (p=0.07 by Welch test), indicating that the dumbbell necks may act as a template where actin networks can grow on membrane domains with their preferred curvature (Fig. 6 D). Both sets of curvatures are also consistent with the curvature of concave actin patches reported by Dürre et al. (20).

## DISCUSSION

To study the interplay of branched actin networks with lipid membranes, we developed a protocol that allows the reproducible and high-yield formation of GUVs with a membrane-nucleated actin cortex. Our method is facile, requires small sample volumes, and is fast enough to prevent premature actin polymerization during encapsulation. It is furthermore compatible with a range of widely used lipid compositions (Fig. S11) and different types and concentrations of density gradient media (Fig. S12 A). We note that commonly used density gradient media such as Optiprep and sucrose influence actin polymerization kinetics (Fig. S12 B-E), highlighting the need for careful experimental design and reporting when encapsulating cytoskeletal polymers.

The new method allowed us to systematically study the formation of actin cortices in GUVs as a function of the concentration of actin and the density of membrane-bound nucleators. We found that Arp2/3-nucleated cortices can grow in different regimes, depending on the amount of actin which is available in the GUV. This allows us to reconcile previous works, which have found that actin forms disconnected patches at low concentrations (3 μM (20)) but continuous actin shells at high concentrations (6.5 μM (25)). This is consistent with the known autocatalytic nucleation mechanism of actin by Arp2/3, which should produce isolated patches: New actin filaments will preferentially be assembled where Arp2/3 can be activated by nucleation promoting factors (71), i.e., at the membrane, and where an existing actin filament is available to act as a primer (8), i.e., in an existing patch. At low density or short filament length (promoted by capping protein), multiple primers generate independent networks. With increasing actin density and/or filament length, these independent networks can merge into a continuous cortex made of entangled filaments. Irrespective of the actin conditions, we found variable phenotypes among GUVs, likely due to a significant variability in protein encapsulation efficiency, which we demonstrated directly by quantitative microscopy. We note that to form minimal actin cortices, the presence of additional regulatory proteins used in earlier works (profilin, gelsolin, ADF/cofilin and capping protein (20, 25)) was not required.

Under conditions where the actin cortex formed distinct patches, we found that the patches generated negatively curved membrane sites. Thus, actin, membrane-bound VCA and Arp2/3 are sufficient to allow for curvature generation by actin on a network scale. Conversely, we found that these minimal dendritic actin networks also recognized membrane curvature: in GUV samples prepared by eDICE, we always observed some dumbbell-shaped GUVs where actin preferentially localized to the neck. These dumbbell GUVs form by a mechanism involving the hemifusion of two nested GUVs, whose combined shape is determined by an interplay of osmotic pressure across the membrane, and line tension in the hemifusion stalk. We found a striking match between the typical membrane curvature generated by concave actin patches in spherical GUVs, and the curvature of the membrane necks of dumbbell GUVs at which preferential network assembly occurs, on the order of 0.5 *μ*m^− 1^. These curvatures qualitatively match the curvature of concave actin patches reported in a previous reconstitution study(20). Thus, actin, membrane-bound VCA and Arp2/3 are sufficient to allow for curvature sensing and generation of actin on a network scale, requiring none of the curvature generating or -sensing binding partners which are thought to mediate curvature sensitivity of actin networks in cells (14–19).

What exactly mediates this network-level curvature sensing remains elusive. It has been suggested that the competition of Arp2/3 and capping protein for actin monomers may be a driving factor behind the growth of curved actin networks (20), although it remains unclear how this mechanism would lead to membrane curvature generation. We found that capping protein was actually not strictly necessary for curvature generation by actin, and that curvature sensing happened robustly in dumbbell-shaped GUVs as long as actin was nucleated on the membrane by Arp2/3. The effect was reproducible across different surface densities of nucleators and occured for different Arp2/3 isoforms.

On a molecular scale, the Arp2/3 complex itself has been shown to preferentially bind to actin filaments on their convex side, i.e., binding sites from which the mother filament bends away, are preferred (72). This effect has been suggested to be amplified on the network level (72), which may help cells orient polymerization forces effectively in organelles like the lamellipodium. Curiously, if preferential assembly of actin on convex mother filaments occurs on an inwardly bent piece of membrane, we should expect curvature preference of Arp2/3 to drive nucleation of the densest networks at the sides of the bent membrane patch, rather than in its middle. If we imagine an initially flat membrane decorated with a branched cortex, where all the filament ends are anchored firmly on the membrane, local inward bending should splay apart the actin branches at the sides of this deformation (thus increasing nucleation there), while branches would be unaffected or slightly compressed at the tip of the deformation, hence lowering the nucleation rate there. By contrast, we observed that actin intensity peaked at the center of the concave membrane patches, which suggests that a different mechanism is behind the formation of concave membrane patches. We note also that our data does not allow us to distinguish whether actin polymerizes on the membrane and subsequently accumulates at dumbbell necks by lateral diffusion, or whether net actin assembly is favoured where the membrane is bent. This distinction would shed light on the mechanism of curvature sensing and generation in branched actin networks, and should be followed up on in future works.

Membrane curvature sensing of actin in the absence of curvature-sensitive proteins to mediate actin-membrane interactions has so far, to the best of our knowledge, never been observed. We speculate that it may present a mechanism by which actin assembly can be robustly directed to actin-rich membrane structures with a micron-scale curvature, such as the early cytokinetic furrow (73), phagocytic cups (11), or during cell migration through physical constrictions that mediate micron-scale inward membrane curvature. Actin enrichment at curved membrane sites has for instance been reported around the nucleus of immune cells squeezing through micrometric constrictions (12), osteoblasts in nanogrooves (74), and fibroblasts migrating on nanopatterned substrates with structure sizes of a few 100 nm in size (75). While other proteins certainly contribute to this organization in live cells, our work raises the question whether an intrinsic curvature sensitivity of branched actin networks may help in making these associations more robust.

Further, the curvature preference of branched actin networks may potentially be used as a design strategy for building synthetic cells with an actin-based division machinery. Researchers in the growing field of bottom-up synthetic biology are pursuing different strategies to promote membrane constriction based on imposing membrane curvature by microfluidic trapping (76, 77) or addition of membrane-shaping proteins (78). Our findings suggest that preferential assembly of actin cortices can be promoted in simpler systems, requiring fewer proteins and thus less careful tuning of encapsulation stoichiometries or *in-vesiculo* transcription and translation.

## CONCLUSION

We developed a robust protocol for actin encapsulation in cell-like lipid containers. We used these actin-containing membrane vesicles to show that dendritic actin networks are able to recognize as well as generate membrane curvature. We observed that cortical actin networks preferentially assembled at the necks of dumbbell-shaped GUVs and were able to form concave membrane patches in spherical GUVs. This finding raises the intriguing possibility that intrinsic curvature sensitivity of branched actin nucleation may contribute to the localization of actin at curved membrane regions in cells such as the early cytokinetic furrow.

## AUTHOR CONTRIBUTIONS

All authors contributed to the research design. L.B. and M.A.P. performed all the experiments and analyzed the data. F.F. performed all the theoretical modeling. L.B. wrote the manuscript together with F.F. All authors reviewed, approved, and contributed to the final version of the manuscript.

## ACKNOWLEDGEMENT

We thank Jeffrey den Haan for protein purification, Kristina Ganzinger (AMOLF) for providing the 10xHis VCA construct, David Kovar (University of Chicago) for the capping protein constructs and Michael Way (Crick Institute) for providing purified human Arp2/3 proteins. We are grateful to Iris Lambert for early actin encapsulation experiments that formed the basis for establishing the eDICE method, to Federico Fanalista for acquiring images of dumbbell-shaped GUVs in samples produced by cDICE, and to Tom Aarts for images of dumbbell shaped GUVs produced by gel-assisted swelling. Lennard van Buren is thanked for his help with image analysis to quantify actin concentrations in GUVs. We thank Kristina Ganzinger (AMOLF) for hosting us to perform pyrene assays in her lab, and Balász Antalicz (AMOLF) for technical assistance with the spectrophotometer. The authors also thank Matthieu Piel and Daniel Fletcher for insightful and inspiring discussions. We acknowledge financial support from The Netherlands Organization of Scientific Research (NWO/OCW) Gravitation program Building a Synthetic Cell (BaSyC) (024.003.019) and from the Kavli Institute of Nanoscience Delft. F.F. gratefully acknowledges funding from the Kavli Synergy program of the Kavli Institute of Nanoscience Delft.

## CODE AVAILABILITY

A python script to generate the predicted dumbbell shapes from our model, as well as Jupyter notebooks to analyze the experimental data, are available on GitHub: https://github.com/BioSoftMatterGroup/actin-curvature-sensing

## SUPPORTING CITATIONS

References (55, 64, 67–70, 79–82) appear in the Supporting Material.

