## Supplemental Information for "Reconstituted branched actin networks sense and generate micron-scale membrane curvature"

### I. THEORETICAL MODEL OF EQUILIBRIUM SHAPES OF DUMBBELL VESICLES

As described in the main text, we assume that a dumbbell GUV is created of two vesicles, a larger outer one containing a smaller inner one, of which the inner one has burst, resulting in a shape with two lobes: one with a single and the other with a double membrane bilayer. Here we study the shape of these dumbbells. Our approach is motivated by earlier theoretical work on domain-induced budding of vesicles composed of phase-separating lipid mixtures [1–4]. First we explain why the dumbbell shape can be described by two spherical caps, and second we explain how we infer the ratio between line tension and vesicle pressure from the shape of the GUV.

The shape energy of a GUV with negligible bending energy is given by [2]:

$$\mathcal{H} = \oint_{\partial 1} \sigma \, \mathrm{d}l + \Sigma^{(1)} A^{(1)} + \Sigma^{(2)} A^{(2)} + PV \, . \tag{1}$$

The first term in Eq. (1) describes the energy associated with the perimeter of the interface between the double bilayer surface and the single bilayer surface. Therefore, the integral runs along the edge of this interface, where $\sigma$ is the line tension. The second and the third term correspond to the membrane tension, where $A_i$ is the surface area and $\Sigma^{(i)}$ is the membrane tension of domain $i$, with $i = 1$ for the double membrane and $i = 2$ for the single membrane domain. The fourth term introduces a volume constraint, where $P$ is the pressure difference between the inside and the outside of the GUV and $V$ is the volume of the GUV.

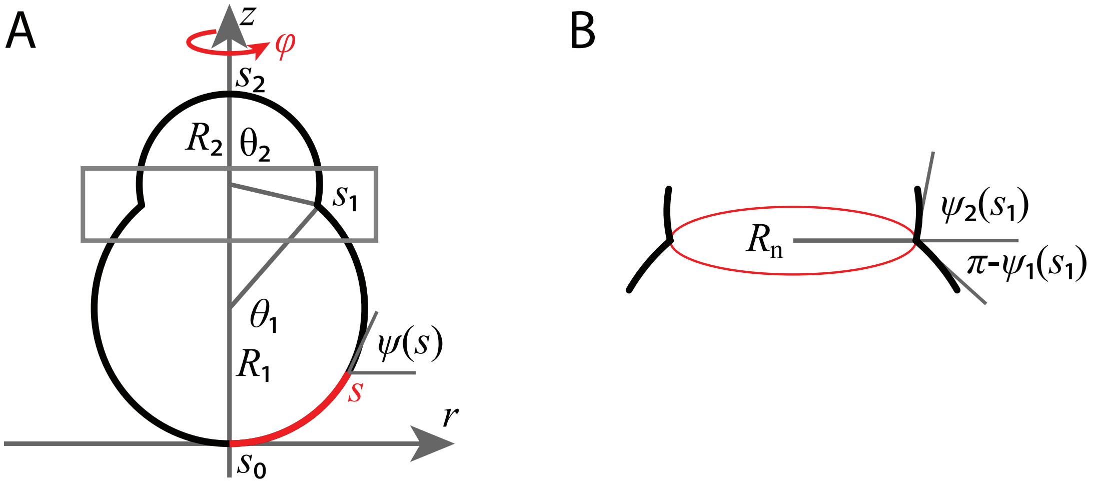


FIG. S1. **Geometry of dumbbell GUVs.** (A) Parametrization of the dumbbell shape by the arc length $s$ (red) along the (rotationally symmetric) membrane contour. We take $s_0 = 0$ at the south pole of the dumbbell. The local orientation of the membrane is given by the angle $\psi(s)$. The bright and dim lobes are spherical caps characterized by their radii $R_1$ and $R_2$ and their opening angles at the neck, $\theta_1$ and $\theta_2$. The neck is located at $s = s_1$, and the north pole at $s = s_2$. (B) Zoom-in to the dumbbell neck region (grey rectangle, panel A). The line tension of the hemifusion line (red) determines the neck radius $R_n$. Dumbbell lobes connect to the plane of the neck at angles $\pi - \psi_1$ and $\psi_2$, respectively.

Importantly, we here consider the limiting case where the energy associated with the line tension and the volume constraint dominates the system, and thus the contribution of the bending energy is negligible in Eq. (1). Since dumbbell shapes are axisymmetric, we can parameterize them by a (trivial) rotation angle $\phi$ about the symmetry axis, and a contour length $s$ along their shape, see Fig. S1. The shape and local orientation of the contour can then be described by the radial position $r(s)$ and the tangent angle $\Psi(s)$. These parameters satisfy the geometric constraint $\dot{r} = \cos\Psi$, with a dot denoting the derivative with respect to the arc length $s$. To enforce this constraint, we add a term $\gamma(\dot{r} - \cos\Psi)$ to the shape energy, where $\gamma$ is a Lagrange multiplier. We can then write the shape energy as the integral over an energy density

$$\begin{aligned}\mathcal{H} = \oint_{\partial 1} \sigma \, \mathrm{d}l &+ 2\pi \int_{s_0}^{s_1} \left[ \Sigma^{(1)} r(s) + \frac{1}{2} P r^2 \sin\psi(s) + \gamma \left( \dot{r} - \cos\psi(s) \right) \right] \mathrm{d}s \\ &+ 2\pi \int_{s_1}^{s_2} \left[ \Sigma^{(2)} r(s) + \frac{1}{2} P r^2 \sin\psi(s) + \gamma \left( \dot{r} - \cos\psi(s) \right) \right] \mathrm{d}s. \end{aligned} \tag{2}$$

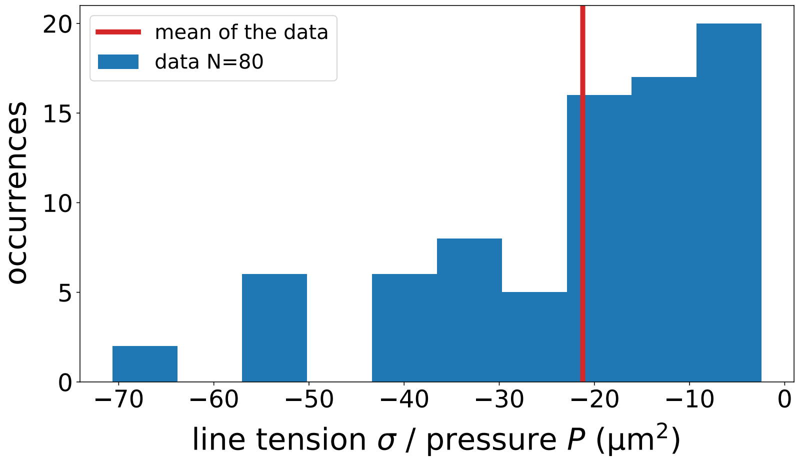


FIG. S2. Histogram of the line tension $\sigma$ divided by the GUV pressure difference $P$ calculated from the measured GUV dumbbell shapes (main text Fig. 4 G,H), according to Eq. (7).

We can minimize the energy density within each domain, which gives us the following shape equations [2]:

$$\begin{aligned} 0 &= \frac{1}{2}Pr^2\cos\psi + \gamma\sin\psi, \\ \dot{\gamma} &= \Sigma^{(i)} + Pr\sin\psi, \\ \dot{r} &= \cos\psi. \end{aligned} \tag{3}$$

Moverover, we get a matching condition at the boundary ($s = s_1$), where there is a jump in the tangent angle $\psi$:

$$\lim_{\varepsilon\downarrow 0}\left[\gamma(s_1+\varepsilon) - \gamma(s_1-\varepsilon)\right] = \sigma. \tag{4}$$

The spherical caps satisfy $\psi(s) = s/R_i$ and $r(s) = R_i\sin(s/R_i)$. As can be easily checked, these spherical caps are indeed solutions of Eqs. (3) [4], if the following relations hold:

$$p = -\frac{2\Sigma^{(i)}}{R_i}, \tag{5}$$

$$\gamma = -\frac{1}{2}pR_i^2\sin\psi\cos\psi = \sigma^{(i)}R_i\sin\psi\cos\psi \tag{6}$$

Eq. (5) is the well-known Laplace pressure, relating the pressure difference across the membrane to the membrane's surface tension. Eq. (6) gives us the value of the Lagrange multiplier $\gamma$ in each domain. The two domains meet at the neck, where $R_1\sin\theta_1 = R_2\sin\theta_2 = R_\mathrm{n}$. Using the expressions for $\gamma$ in the matching condition, Eq. (4), with $\psi_i = \theta_1$ and $\psi_2 = \pi - \theta_2$, allows us to calculate the ratio between the line tension $\sigma$ and the pressure gradient $P$ [3, 5]:

$$\frac{\sigma}{P} = \frac{1}{2}R_\mathrm{n}^2\left(\cot\psi_1 - \cot\psi_2\right) = \frac{1}{2}R_\mathrm{n}^2\left(\cot\theta_1 + \cot\theta_2\right) \tag{7}$$

Fig. S2 shows the histogram of the ratio $\sigma/P$ (blue) calculated from the measured GUV dumbbell shapes (cf. main text Fig. 4 G,H), according to Eq. (7). Here, the opening angles of the bright and dark lobe of the dumbbell GUV, $\theta_1$ and $\theta_2$, are calculated from the diameters of the bright and dark lobe and the neck diameter of the GUV. The mean of the histogram (red) is at $-21\,\mu\mathrm{m}^2$, which is about half of the measured value for a vesicle that undergoes domain-induced budding ($-49\,\mu\mathrm{m}^2$) [3].

#### II. SUPPLEMENTAL EXPERIMENTAL METHODS

##### A. Supplemental materials

Cholesterol (3$\beta$-Hydroxy-5-cholestene, 5-Cholesten-3$\beta$-ol, Cat. # C8667-1G) and sucrose (Cat. # S0389) were purchased from Sigma Aldrich. The following lipids were purchased from Avanti Polar Lipids: L-$\alpha$-phosphatidylcholine

(95 %) from chicken egg (EggPC), 1,2-dioleoyl-sn-glycero-3-phospho-L-serine (DOPS), 1,2-dioleoyl-sn-glycero-3-phospho-(1'-myo-inositol-4',5'-bisphosphate) (ammonium salt) ($PIP_2$), 1',3'-bis[1,2-dioleoyl-sn-glycero-3-phospho]-glycerol (sodium salt) (cardiolipin), and 1,2-dioleoyl-sn-glycero-3-phosphoethanolamine-N-(cap biotinyl) (biotin-PE). $PIP_2$ was stored in a mixture of chloroform, methanol and water at a 20:9:1 volumetric ratio, all other lipids were stored in chloroform and under argon at -20° C.

##### B. Supplemental microscopy methods

Widefield microscopy images were acquired on an inverted Leica Thunder Imager widefield microscope equipped with a 200 mW solid state LED5 light source, a 63x water immersion microscope (HC PL APO 63x / 1.20 W Corr CS2) and a monochrome sCMOS camera (Leica). Epifluorescence images were acquired on an inverted Nikon Ti Eclipse microscope equipped with a 60x water immersion objective (CFI Plan Apochromat VC), a digital CMOS camera (Orca Flash 4.0), and an LED light source (Lumencor Spectra Pad X). Phase contrast images were acquired on the same Nikon Ti microscope using its DIA illuminator at a voltage of 12 V and using the corresponding phase mask in the microscope's condenser.

##### C. GUV production by gel-assisted swelling

Gel-assisted swelling of GUVs was performed following Ref. [6]. Cover glasses (22 x 22 mm, No. 1.5H, Paul Marienfield GmbH & Co. KG) were first rinsed with ethanol and MilliQ water and dried under a stream of nitrogen. They were then plasma cleaned for 30 seconds (PlasmaPrep III, SPI supplies), after which 100 $\mu$L of a solution of 5 % (w/v) polyvinyl alcohol (PVA, 145 kDA, 98 % hydrolysed, VWR, Amsterdam the Netherlands) in 200 mM sucrose in MilliQ water was spread over each coverslip at room temperature. The gel was solidified by baking it in an oven for 30 minutes at 50 ° C. Then, 10 $\mu$L of a lipid solution at a total lipid concentration of 1 mg/mL in chloroform, consisting of DOPC:Atto488-DOPE at a molar ratio of 99.5:0.5, was spread over the gel. The gel was placed in a vacuum desiccator for 30 minutes to ensure total evaporation of the organic solvent. The cover glasses were then placed in a compartmentalized petri dish (4 compartments, VWR), and to each gel we gently added 300 $\mu$L of GUV swelling buffer (10 mM Tris-HCl at pH 7.4, 100 mM KCl, 100 mOsm sucrose). After swelling for one hour, GUVs were collected by taking up the swelling solution with a pipette (1 mL tip), flushing the solution again over the cover slip once to dislodge the GUVs, and pipetting it up again.

##### D. GUV production by cDICE

cDICE GUVs were prepared from the same solutions and in the same rotating chamber as eDICE GUVs. Droplets were formed by injecting IAS into the oil phase at a rate of 25 $\mu$L/min for 5 min using a syringe pump (KDS 100 CE, KD Scientific). The capillary setup was identical to that described in [7].

##### E. Actin polymerization assays

Actin polymerization was quantified by the classical pyrene actin assay [8], following an existing protocol [9], briefly summarized below.

*a. Hardware* Pyrene assays were performed on a Duetta fluorescence and absorbance spectrometer (Horiba Scientific), equipped with an 80 W S/N 1344-DL lamp and TC1 temperature controller (Quantum Northwest). The temperature varied between 25.0 and 25.4 ° C. Measurements were performed following a custom-defined procedure in the Horiba EzSpec software. For an excitation wavelength of 365 nm, the emission at 407 nm was recorded for a 1 s integration time per data point. The excitation window was set at 10 nm and the emission window at 5 nm. These parameters were chosen to maximise the relative difference in intensity between polymerized and unpolymerized pyrene-actin samples. All samples were measured in 3-window Quarz microvolume cuvettes from Hellma Analytics with an optical path length of 3 mm, holding 55 $\mu$L sample volumes. Cuvettes were cleaned by rinsing with 5 mL MilliQ water, 2 mL ethanol, and 2 mL MilliQ water between measurements, and dried under $N_2$ stream. Between different days of measurements, the cuvettes were further cleaned by sonicating in a 1 % Hellmanex solution for 5 min, sonicating for 5 min in MilliQ water to remove any excess detergent, and finally rinsing with 5 mL MilliQ water before drying under $N_2$ stream.

*b. Procedure* We performed all experiments in a final buffer with the same composition as the GUV IAS buffer, with at least two independent measurements per condition. Measurements with the different Arp2/3 isoforms but no optiprep in the buffer were performed once per isoform. The final concentrations of actin, VCA and Arp2/3 complex were 4 $\mu$M, 650 nM, and 50 nM, respectively. We first prepared an actin premix to ensure a consistent starting concentration and labeling ratio of monomeric actin. It contained 95 % unmodified actin and 5 % pyrene-labeled actin, at 23.8 $\mu$M total, corresponding to 5.95 x the final actin concentration for the pyrene assay, in G-buffer. In the cuvette, the actin premix was then diluted into Mg-G-buffer (5 mM TrisHCl pH 7.4, 0.2 mM $MgCl_2$, 1 mM DTT, and 0.2 mM MgATP) and allowed to incubate for 2 min. In this step, $Ca^{2+}$ ions on actin monomers are exchanged for $Mg^{2+}$ ions, ensuring that actin polymerization commences at a reproducible starting point. Salts, Arp2/3 and VCA were then added and mixed by pipetting 5 times. The cuvette was placed in the spectrophotometer and the measurement was started, noting down the delay between adding salt and the first recorded data point (usually 10-15 s). The measurement was left to run until the fluorescence intensity plateaued.

*c. Data analysis* While the Duetta spectrometer offers simultaneous buffer measurement and correction, we chose to perform a manual background correction to increase time resolution of our measurements. This allowed us to acquire one datapoint per 1.1 s. For every measurement day, we first performed blank measurements on F-buffer and F-buffer including 6.5 % optiprep, where we repeated the same measurement 20 times. We then averaged these data points for every buffer individually, and subtracted the average from the sample data before further processing. To extract the time it takes to reach maximum pyrene fluorescence, as well as the actin elongation rate at the time where half of all G-actin is polymerized, we use a custom-written python script following the analysis laid out in [9]. We extract the time it takes each sample to reach maximum pyrene fluorescence and hence steady-state actin polymerization, as well as the actin elongation speed at the inflection point of the pyrene fluorescence curve. At this point, it is assumed half of all actin has polymerized and the increase in fluorescence intensity is only due to elongation, not nucleation [9].

### III. SUPPLEMENTAL FIGURES

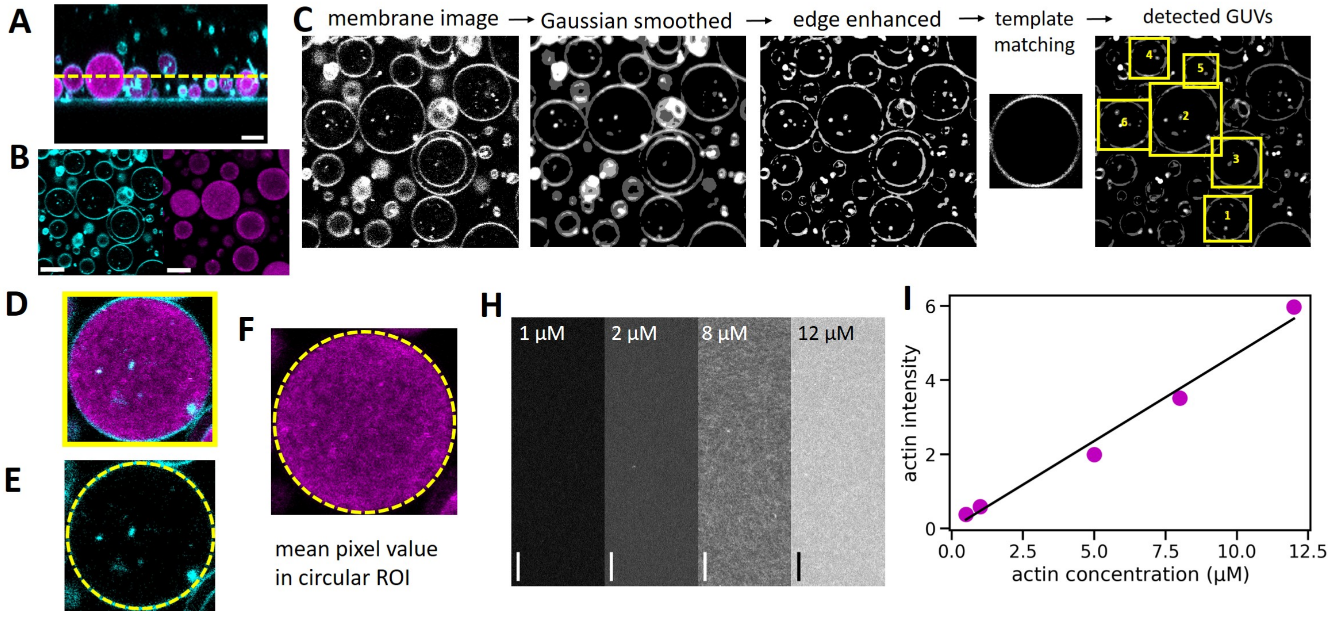


FIG. S3. **Quantifying actin concentrations.** (A) For quantitative confocal imaging, we performed an xz-scan to locate the precise z-position of the coverslip visible from its reflection (horizontal cyan line). Actin is shown in magenta, lipids in cyan. The yellow dashed line indicates a 7 $\mu$m height above the coverslip, where we took subsequent images. (B) Next, we acquired a two-channel xy-confocal image (actin in magenta, lipids in cyan). (C) We used an automated pipeline to detect the vesicles by pre-processing the images and subsequently locating GUVs using the template matching module in the DisGUVery toolbox [10]. Yellow boxes indicate the detected GUVs. (D) To extract the actin intensity inside a given GUV, we used the original unprocessed two-color image. We loaded each GUV detected by the template matching algorithm (here: GUV 2 from panel C) and drew a circular ROI of the same size as the template matching box ((E), yellow dashed circle). The mean pixel value inside this circular ROI in the actin channel was then taken as the average actin intensity of the GUV. To convert actin intensities into concentrations, we acquired a series of reference images of bulk actin networks (H) and constructed a concentration-intensity-calibration curve (I). Magenta datapoints represent the average mean pixel values of at least four 2048x2048 px reference images per condition, with standard deviations smaller than the data points. We fit the datapoints with a proportional model ($I_{\mathrm{act}}(c_{\mathrm{act}}) = A \cdot c_{\mathrm{act}}$, black line), which yields a proportionality constant of 0.471. Scale bars: 5 $\mu$m.

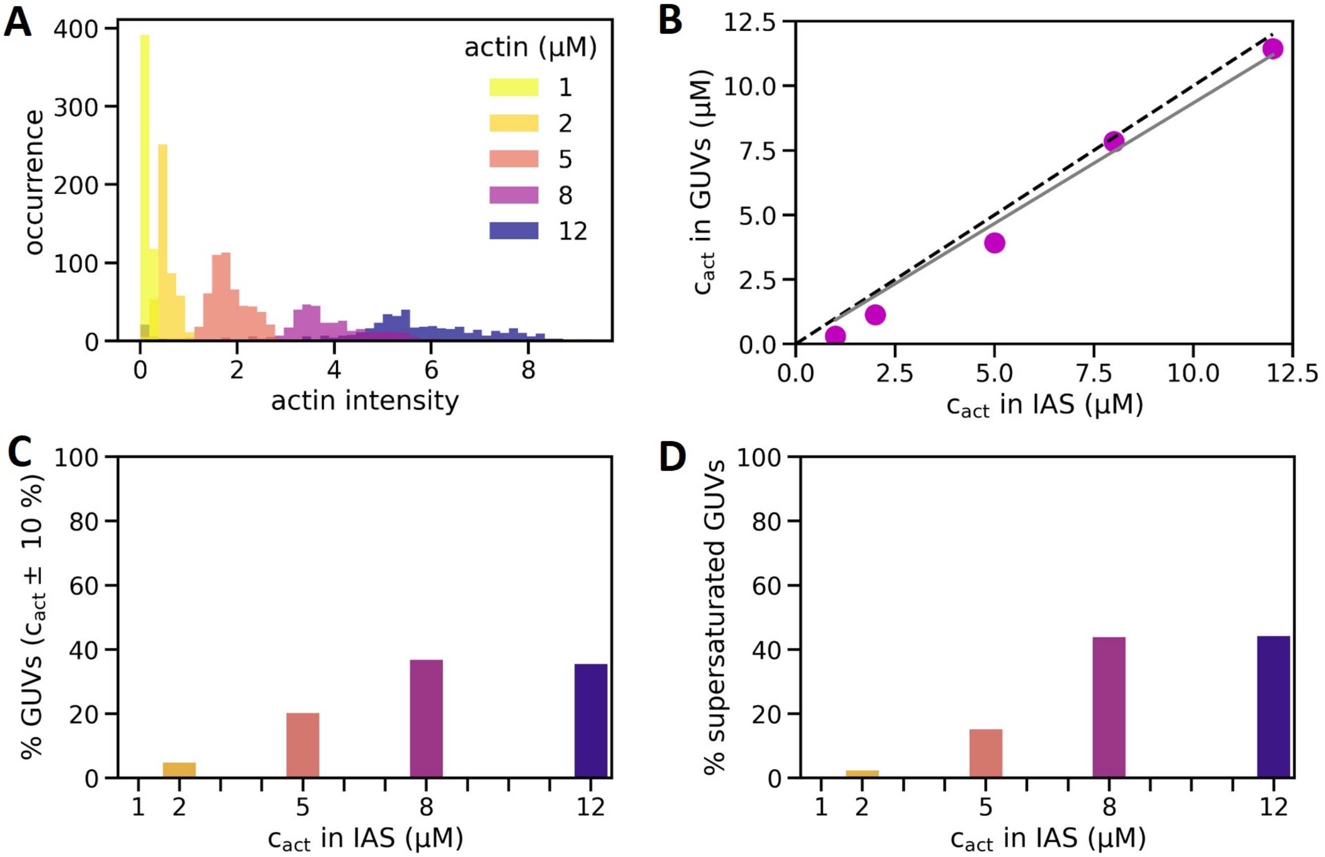


FIG. S4. **Extended quantification of actin encapsulation efficiency by eDICE.** (A) Histogram of actin intensities measured in individual GUVs by quantitative confocal microscopy. Darker colours indicate higher nominal actin concentrations. N = 2334 GUVs, with individual sample sizes $N_i$ = 515, 468, 536, 368 and 447 GUVs for i = 1, 2, 5, 8, and 12 $\mu$M actin, respectively. (B) Mean actin concentration in the GUVs as a function of the nominal actin concentration in the inner aqueous solution. Magenta datapoints represent the mean of all GUVs at one nominal concentration (same sample sizes as in (A)), and the solid grey line indicates the proportional fit $c_{\mathrm{GUV}}(c_{\mathrm{IAS}}) = \xi \cdot c_{\mathrm{IAS}}$, with $\xi = 93.3\%$. The dashed black line indicates perfect encapsulation, where the concentration in the GUVs equals the input concentration in the IAS. Error bars (95 % confidence interval on the mean) are not shown since they are smaller than the data points. (C) Bar plot showing the fraction of GUVs at different nominal actin concentrations containing actin at approximately the same concentration as the IAS (concentrations at $c_{\mathrm{nominal}} \pm 10$ %). (D) Bar plot showing the fraction of GUVs that are significantly supersaturated in actin ($c_{\mathrm{GUV}} > 1.1 \cdot c_{\mathrm{nominal}}$).

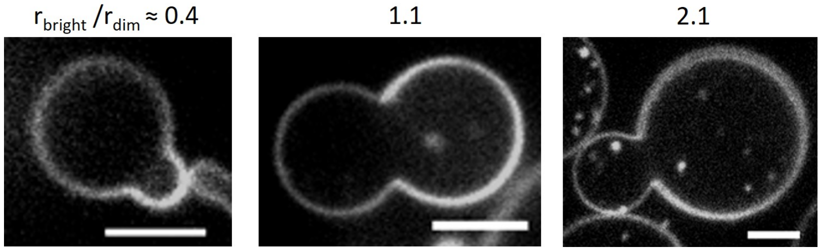


FIG. S5. **Dumbbell GUVs exhibit a range of shapes.** Most dumbbells had nearly equal sized lobes (middel panel, $r_{\mathrm{bright}}/r_{\mathrm{dim}} \approx 1$), but both significantly lower and higher size ratios also occurred (left and right panel, $r_{\mathrm{bright}}/r_{\mathrm{dim}} \approx 0.4$ and 2.1, respectively).

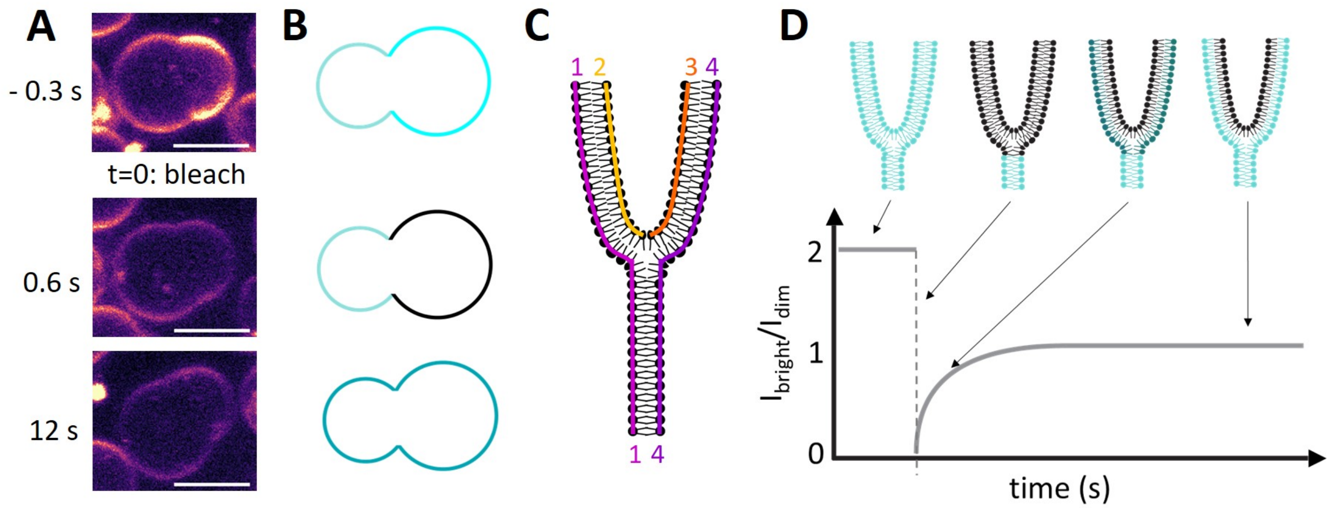


FIG. S6. **FRAP analysis to test a mechanism for dumbbell formation based on hemifusion of nested GUVs.** (A) Confocal images taken at the equatorial plane of a dumbbell-shaped GUV before and after bleaching the bright lobe. Scale bars: 5 $\mu$m. Membrane fluorescence is shown in false color (magma) for clarity. (B) Schematic of the bleaching experiment. The GUV starts out with a dim and a bright lobe (top). Ideally, all of the bright lobe's fluorescence gets bleached (middle). The remaining fluorescence from the unbleached lobe then redistributes in all connected leaflets of both lobes. (C) Schematic of the different lipid leaflets at the neck, assuming that the dumbbell shape results from hemifusion: leaflets 1 and 4 connect the dim and bright lobe, while leaflets 2 and 3 are disconnected and only span the bright lobe. Note that 'leaflets' 2 and 3 are two halves of one continuous monolayer of lipids that is bent in on itself, making it macroscopically appear as two separate leaflets. (D) Expected ideal recovery curve (bottom) and illustration of the fluorescence distribution in the different leaflets (top). Leaflets 2 and 3 (C) do not exchange lipids with leaflets 1 and 4, and consequently remain dark after bleaching. The intensity ratio between the bright and dim dumbbell lobes thus recovers not to 2, but to 1.

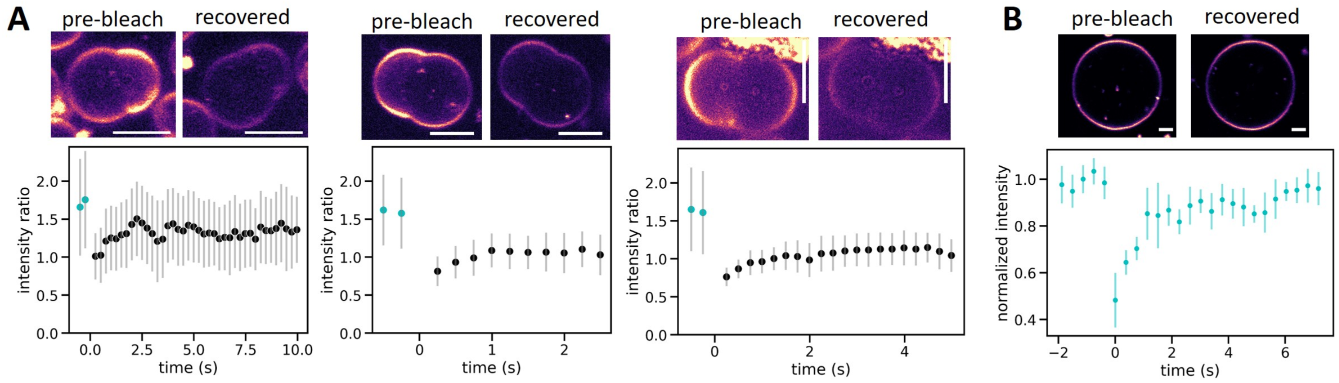


FIG. S7. **Extended FRAP data for dumbbell-shaped GUVs.** (A) Example FRAP curves for three additional dumbbells not shown in the main text. The bright lobe was bleached to approximately 30 % of its initial fluorescence intensity at t=0. The intensity ratio between the bright and the dim lobe consistently started out at around 1.7 (cyan data points), and recovered to between 1.0 and 1.4 within 2 s (black data points). (B) FRAP curve of a non-dumbbell GUV. Here, a 2x5 $\mu$m rectangular ROI at the top of the GUV was photobleached and the fluorescence recovery was monitored relative to a reference region on the opposite side of the GUV. The effect of photobleaching appeared more pronounced here than in the dumbbell GUVs, as we could bleach the small region much faster and were thus able to capture more of the initial recovery period. The fluorescence recovery timescale matched that seen in dumbbell GUVs (< 3 s). Confocal images above each graph show the equatorial slice of the GUV before photobleaching and after at least 15 s of recovery. Scale bars: 3 $\mu$m. Membrane fluorescence is shown in false color (magma) for clarity.

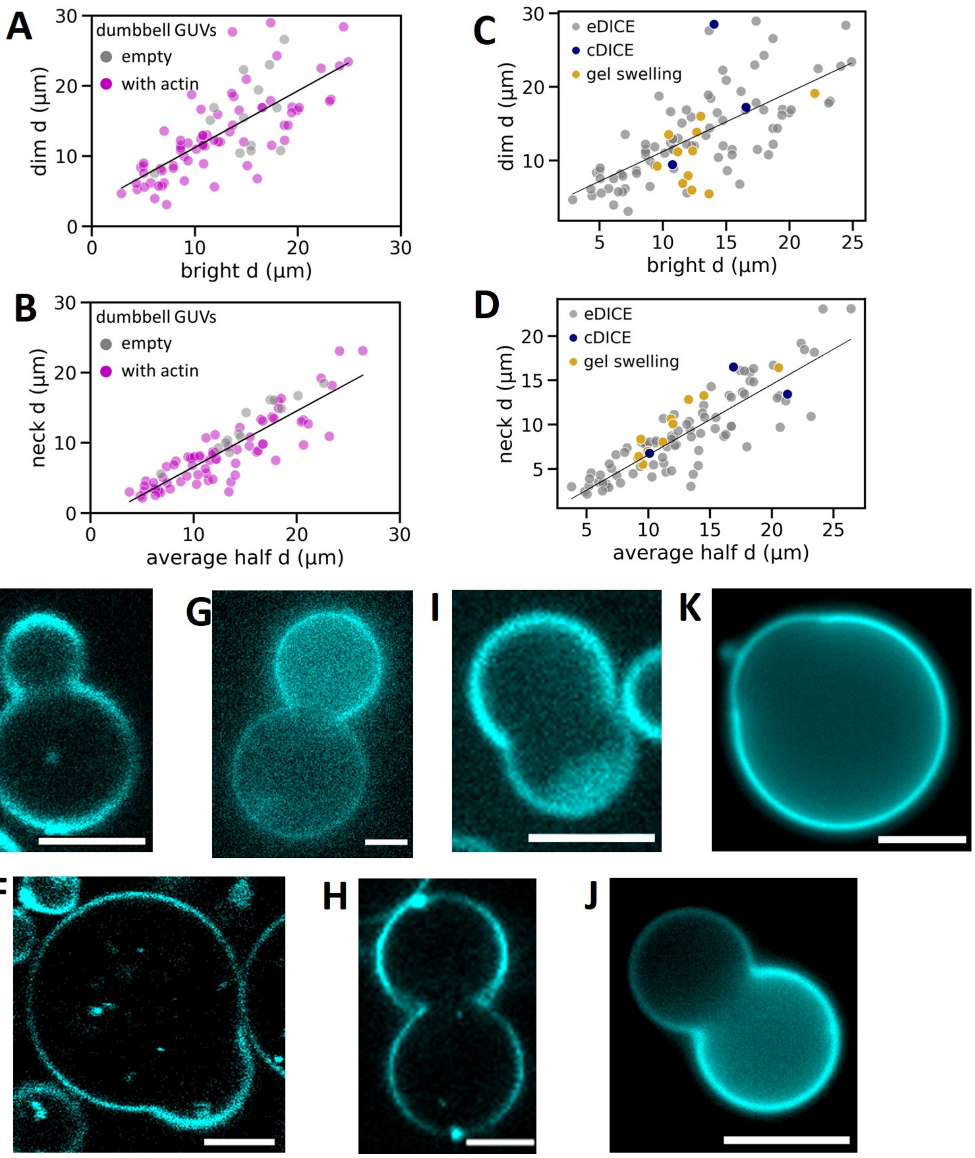


FIG. S8. **Dumbbell GUVs form for a range of IAS compositions and GUV production methods.** (A, B) Data from Fig. 4 H and I are re-plotted to show which dumbbells contain actin (magenta datapoints, either 4.4 or 8 $\mu$M actin in F-buffer, N = 72 GUVs) or only F-buffer (grey datapoints, N = 13). Actin does not appear to alter the formation of dumbbell shaped GUVs. (C, D) The shapes of dumbbells produced by cDICE (blue datapoints, N=3) and gel swelling (yellow datapoints, N=11) follow the same trends in relative lobe dimensions (C) and neck dimensions (D) as those produced by eDICE (grey datapoints, N=72). (E-L) Fluorescence microscopy images of dumbbell shaped GUVs produced in different circumstances. Dumbbells form in GUVs encapsulating only buffer (E), actin which polymerizes in the lumen without nucleating proteins (F), different concentrations of actin nucleated on the membrane by Arp2/3 and VCA (G: 4 $\mu$M actin, H: 8 $\mu$M actin), and with cortices where Arp2/3 driven actin nucleation is modulated by capping protein (I). (J) Dumbbell vesicles are also occasionally observed in samples produced by cDICE, in this case encapsulating G-buffer. (K) We even find dumbbells (albeit only rarely, $< 2$ %) when GUVs are produced by gel-assisted swelling and encapsulate only a sucrose solution (L). E, F, and H were acquired on a laser scanning confocal microscope; G and I were acquired on a spinning disk confocal microscope, and J and K were acquired on a Nikon epifluorescence microscope. Scale bars: 5 $\mu$m.

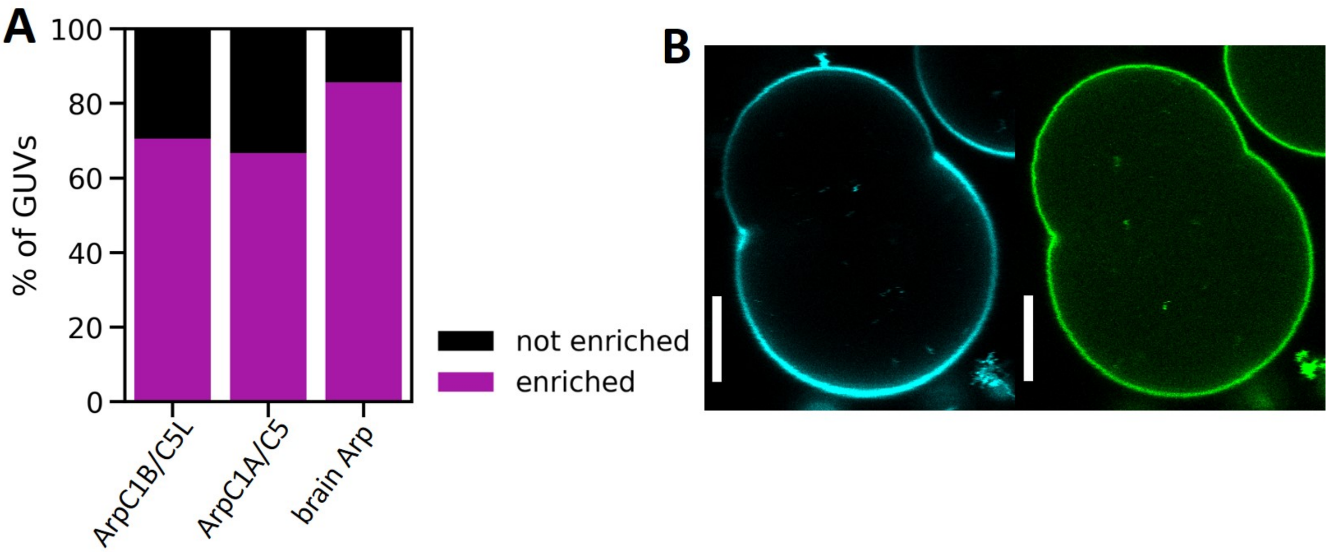


FIG. S9. **Actin was consistently enriched at the necks of dumbbell GUVs for different Arp2/3 isoforms, while VCA alone did not show enrichment.** (A) Bar plot showing the fraction of dumbbell GUVs in which actin was enriched at the neck, depending on the isoform of Arp2/3 which nucleates actin at the cortex. Actin and Arp2/3 were always at 8 $\mu$M and 50 nM, respectively, and VCA was either at 2 or 6.5 $\mu$M. Capping protein was absent. N = 17, 3 and 7 for Arp2/3 isoforms Arp2/3C1BC5L, Arp2/3C1AC5, and mixed isoforms isolated from porcine brain, respectively. While statistics for Arp2/3C1AC5 and brain Arp2/3are low, the data suggest that dendritic actin networks preferentially assemble at concave GUV neck regions for all these Arp2/3 isoforms. (B) Representative confocal image of a dumbbell GUV (membrane in cyan) encapsulating 6.5 $\mu$M fluorescent VCA (green). VCA covered the inner leaflet homogeneously, and we did not observe VCA enrichment at any of the 5 observed dumbbell necks.

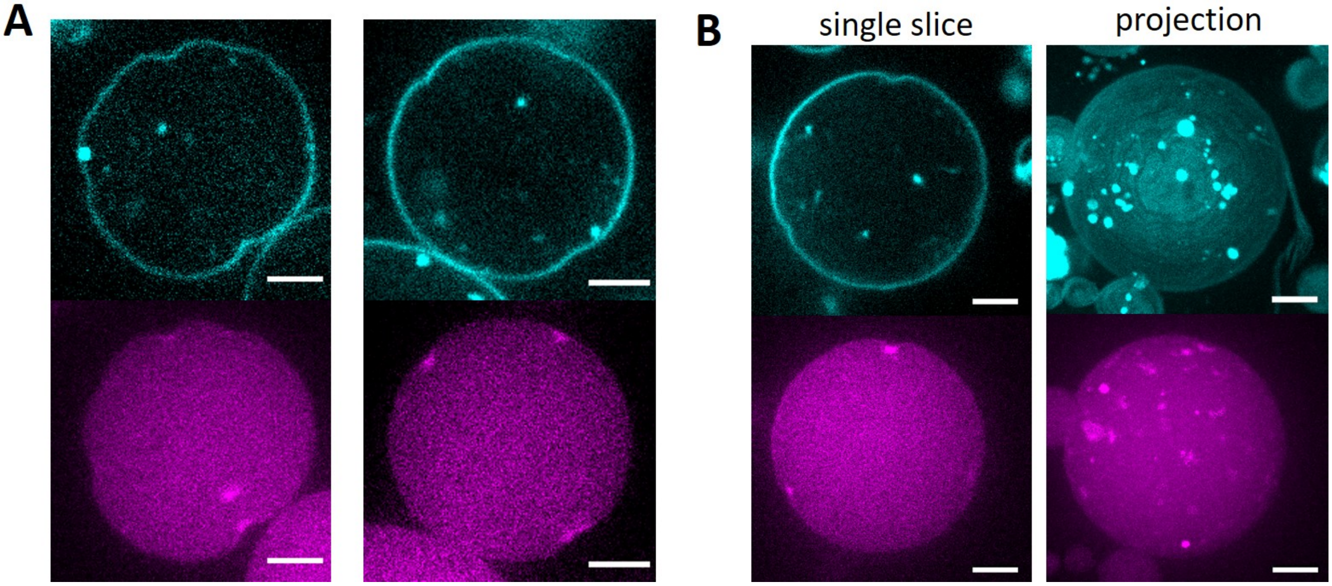


FIG. S10. **Concave actin patches in the presence of capping protein.** Confocal microscopy slices of two representative GUVs containing 8 $\mu$M actin, 50 nM Arp2/3 and 2.6 $\mu$M VCA, combined with 185 nM capping protein. Under these conditions, we observed small regions in which actin (magenta) was enriched at an inwardly bent membrane region (cyan). Typically, GUVs had a few (2-5) patches like the two examples shown in (A), but they could also bear many more concave patches spaced all around the GUV (B). The left and right panels in (B) show a single confocal slice and a maximum intensity projection of the same GUV, and all bright spots visible in the projection represent concave patches. Scale bars: 5 $\mu$m.

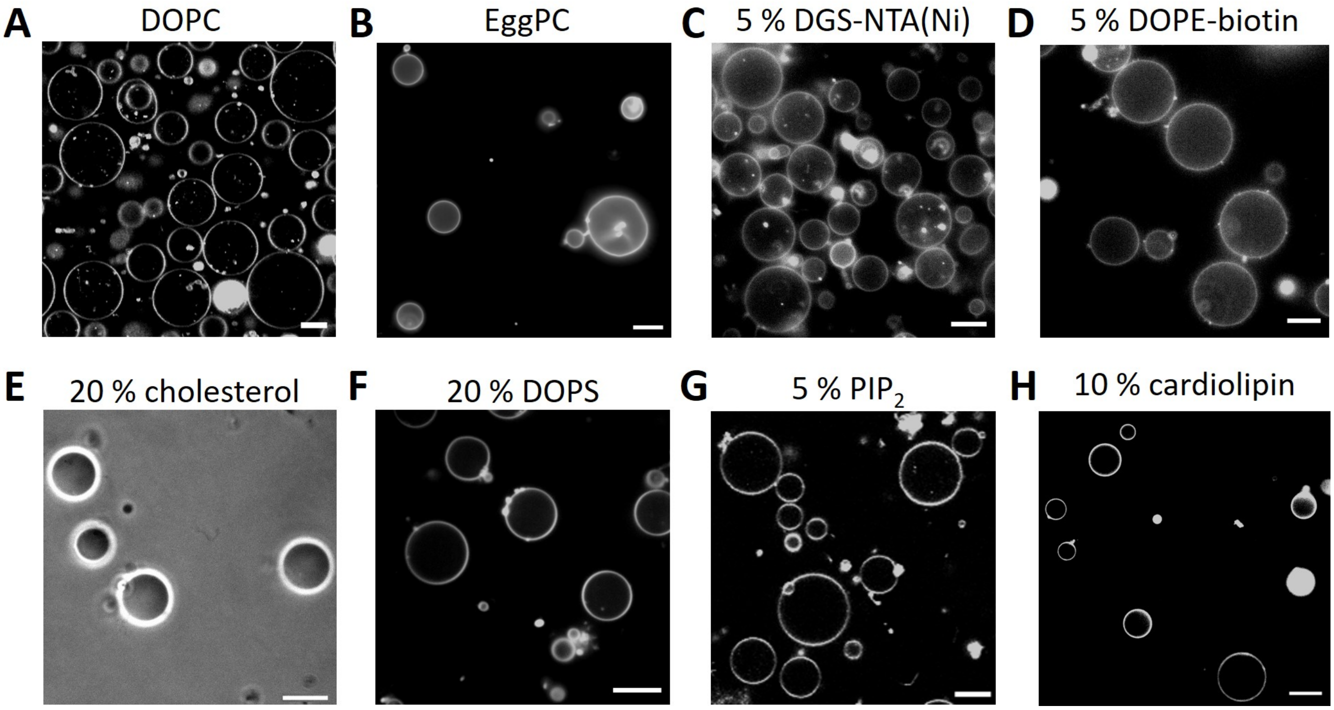


FIG. S11. **eDICE successfully produces GUVs with different lipid compositions.** (A) DOPC, (B) EggPC, (C) 95 % DOPC and 5 % DGS-NTA(Ni), (D) 95 % DOPC and 5 % biotin-DOPE, (E) 80 % DOPC and 20 % cholesterol, (F) 80 % DOPC and 20 % DOPS, (G) 95 % DOPC and 5 % PIP2, (H) 90 % DOPC and 10 % cardiolipin. (A) and (H) are confocal images acquired on the Stellaris LSCM, (B)-(D) and (H) are epifluorescence images acquired on the Nikon epifluorescence microscope, (E) was acquired on the same Nikon microscope in phase contrast mode, and (F) is a confocal image taken on the spinning disk confocal microscope. All GUVs except those shown in (E) are labeled with either 0.005 or 0.05 % of Cy5-DOPE. Scale bars: 10 $\mu$m.

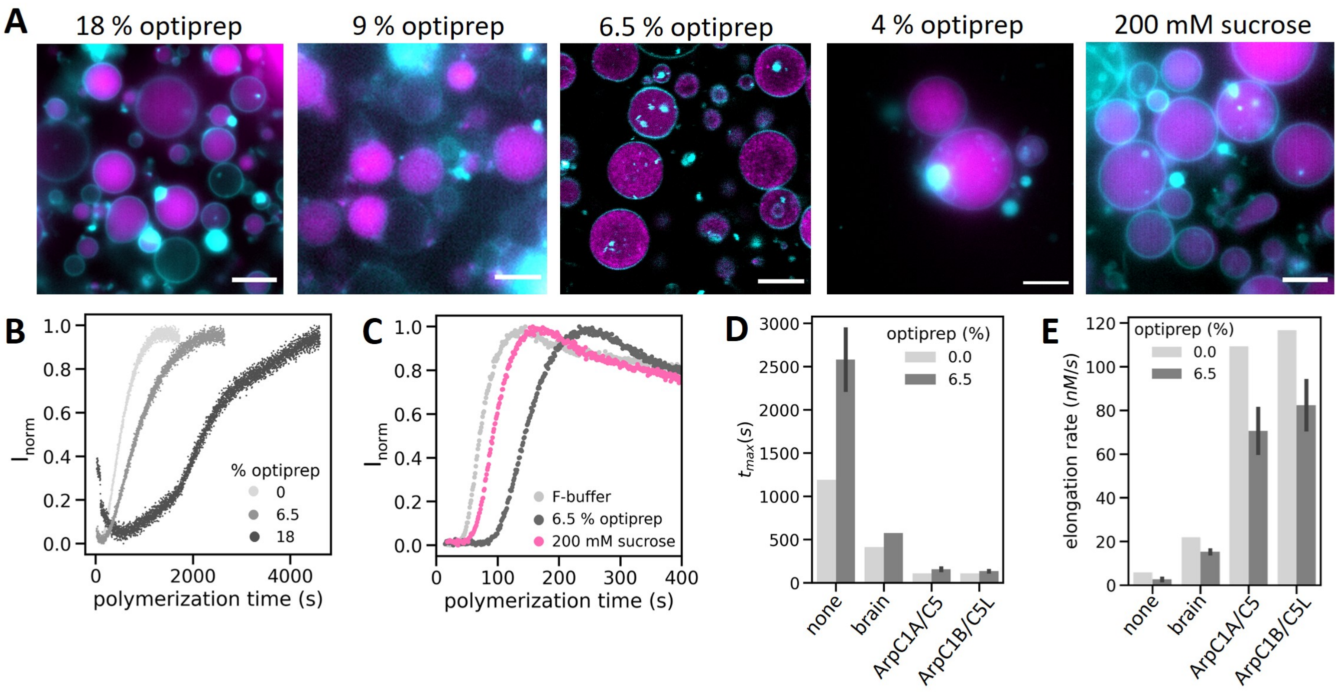


FIG. S12. **Effect of density gradient media on actin encapsulation and polymerization.** (A) Fluorescence images of GUVs encapsulating F-actin using different concentrations of density gradient media (see legends) in the IAS. The GUVs containing 6.5 % optiprep are shown as a single confocal slice, the other images show widefield images. Lipids are shown in cyan, actin in magenta. Scale bars: 10 $\mu$m. (B) Actin polymerization curves measured by pyrene fluorescence assay at different concentrations of Optiprep showed that optiprep interferes significantly with actin polymerization. (C) Pyrene actin polymerization curves indicate that sucrose used at a concentration compatible with eDICE also slowed actin polymerization dynamics, but to a lesser degree than Optiprep. (D) Bar plot of the time to reach steady state, $t_{\max}$ in the presence and absence of 6.5 % optiprep. (E) Bar plot of the actin elongation rate. Light grey bars in panels D,E represent one measurement in F-buffer, and dark grey bars represent the means of at least two separate measurements in F-buffer with optiprep, with error bars indicating the full spread of the data.

#### 98 IV. SUPPLEMENTAL TABLES

| Compound | Instrument | Excitation | Detection |
|---|---|---|---|
| actin-AF488 | Olympus spinning disk | 491 nm laser, 10.5 %, 200 ms | Andor iXon X3 EM-CCD, gain 250 |
| membrane-Cy5 | Olympus spinning disk | 640 nm laser, 53 %, 200 ms | Andor iXon X3 EM-CCD, gain 250 |
| actin-AF488 | Leica Stellaris LSCM | WLL at 499 nm, 18 %, 3.16 µs dwell time | HyDS2 (504-590 nm), counting mode |
| membrane-Cy5 | Leica Stellaris LSCM | WLL at 640 nm, 2 %, 3.16 µs dwell time | HyDX3 (658-809 nm), counting mode |
| VCA-AF488C5 | Leica Stellaris LSCM | WLL at 499 nm, 2 %, 3.16 µs dwell time | HyDX1 (510-567 nm), Standard mode, gain 60 |
| actin-AF488 | Nikon widefield | 470 nm LED, 25 %, 200 ms | Hamamatsu Orca Flash 4.0 |
| membrane-Cy5 | Nikon widefield | 640 nm LED, 30 %, 50 ms | Hamamatsu Orca Flash 4.0 |
| actin-AF488 | Leica widefield | 475 nm LED, 5 %, 200 ms | sCMOS camera, gain 2 |
| membrane-Cy5 | Leica widefield | 635 nm LED, 5 %, 50 ms | sCMOS camera, gain 2 |

TABLE S1. List of imaging settings. The left column lists the compound and their label, which is Alexa Fluor 488 (AF488) or Cy5. The 'Excitation' column shows the excitation wavelength, laser attenuation, and exposure time (camera-based microscopes) or pixel dwell time (confocal scanning microscope).
